# A Subset of Cortical Areas Exhibit Adult-like Functional Network Patterns in Early Childhood

**DOI:** 10.1101/2024.07.31.606025

**Authors:** Jiaxin Cindy Tu, Yu Wang, Xintian Wang, Donna Dierker, Chloe M. Sobolewski, Trevor K. M. Day, Omid Kardan, Óscar Miranda-Domínguez, Lucille A. Moore, Eric Feczko, Damien A. Fair, Jed T. Elison, Evan M. Gordon, Timothy O. Laumann, Adam T. Eggebrecht, Muriah D. Wheelock

**Author notes:** Corresponding Author Muriah D. Wheelock Mallinckrodt Institute of Radiology 4525 Scott Ave St. Louis, MO 63110.

## Abstract

The human cerebral cortex contains groups of areas that support sensory, motor, cognitive, and affective functions, often categorized into functional networks. These networks show stronger internal and weaker external functional connectivity (FC), with FC profiles more similar within the same network. Previous studies have shown these networks develop from nascent forms before birth to their mature, adult-like structures in childhood. However, these analyses often rely on adult functional network definitions. This study assesses the potential misidentification of infant functional networks when using adult models and explores the consequences and possible solutions to this problem.

Our findings suggest that although adult networks only marginally describe infant FC organization better than chance, misidentification is primarily driven by specific areas. Restricting functional networks to areas with adult-like network clustering revealed consistent within-network FC across scans and throughout development. These areas are also near locations with low network identity variability. Our results highlight the implications of using adult networks for infants and offer guidance for selecting and utilizing functional network models based on research questions and scenarios.

**Highlights:**

- Previous studies primarily investigated age-specific networks in infants, with limited focus on how well adult networks describe infant functional connectivity (FC).
- ur analysis identified a subset of areas in infants showing adult-like network organization, where within-network FC shows less age-related variation and higher scan-to-scan reliability.
- These areas are positioned near locations with low variability in functional network identity in adults, indicating a potential link between developmental sequence and interindividual variability in functional network organization.

## 1. Introduction

The human cerebral cortex comprises specialized, large-scale, functional networks (Power et al., 2011; Yeo et al., 2011), supporting sensory, motor, higher-cognitive, and affective functions (Petersen & Sporns, 2015; Wig, 2017). These networks help segregate information processing across different sensory modalities and cognitive domains (Grayson & Fair, 2017; Petersen & Sporns, 2015). In adults, these networks consistently exhibit similar spatial topographies across various acquisition paradigms (task and resting states), and individuals (Gratton et al., 2018). They can also be disrupted by disease (Fornito et al., 2015; Fox & Greicius, 2010). Research indicates that functional networks develop from infancy through old age (Grayson & Fair, 2017; Sun et al., 2023; Wig, 2017), mirroring the progression of complex behavior functions (Grayson & Fair, 2017; Petersen & Sporns, 2015). Preliminary forms of adult functional networks are detectable in utero (Thomason et al., 2013; Turk et al., 2019).

Robust, sometimes bilateral, segregated networks for somatomotor, primary auditory, primary visual, and extrastriate visual cortex are present in infants, whereas higher-order networks appear less mature in early infancy (Eyre et al., 2021; Fransson et al., 2007; Gao, Alcauter, Elton, et al., 2015, 2015; Moore et al., 2024; Myers et al., 2024; Smyser et al., 2010; Sylvester et al., 2022). By 1-2 years, the default mode network becomes more adult-like in some studies (Gao, Alcauter, Elton, et al., 2015; Gao, Alcauter, Smith, et al., 2015; Gao et al., 2009) but remains localized in others (Eggebrecht et al., 2017; Kardan et al., 2022; Marrus et al., 2018; F. Wang et al., 2023).

Researchers analyzing infant neuroimaging data often face a dilemma in choosing an appropriate representation model. Some use the adult network models to describe the relationships between functional connectivity (FC) and behavioral phenotypes in infants (Nielsen et al., 2022; Rudolph et al., 2018; Tooley et al., 2024) or to compare infants and adults (Yates et al., 2023). This choice is sometimes justified by the need to enhance biological interpretability and facilitate communication using consistent terminology across different research groups. However, applying exact adult topography to define functional networks in infants is prone to inaccuracies and may cause the mixing of fMRI BOLD signals (Smith et al., 2011) across different functional networks, reducing statistical power. Additionally, differences in FC may stem from variations in network topography or identity (Bijsterbosch et al., 2018, 2019). Prior-based precision functional mapping using an adult network atlas is also commonly employed in pediatric cohorts (Hermosillo et al., 2024; Moore et al., 2024; Sun et al., 2023). However, these methods assume that infant network organization closely resembles that of adults, which may not be valid.

An alternative approach involves deriving data-driven putative functional networks specific to each developmental stage (Eggebrecht et al., 2017; Kardan et al., 2022; Marrus et al., 2018; Tu et al., 2024; Wheelock et al., 2019). This method could potentially address the issue of poor FC representation within functional networks and improve reproducibility. However, the utility and interpretability of these putative functional networks are less clear, as these putative functional networks have not been validated through task activation studies. Ultimately, the choice of network model should depend on the research goal. It is also crucial to understand the extent to which adult network topography fits infant data. If the adult functional network topography differs significantly from that of infants, using adult functional networks in infant studies could lead to low reliability (Marek et al., 2022). Here, we aim to investigate this issue further and examine the similarities and differences between adult and infant networks in describing the modular structure of FC.

Converging evidence from various modalities indicates that the human cortex develops non-uniformly across different areas. Primary sensory and motor cortex areas mature earlier than higher-order association cortex areas (Ahmad et al., 2023; Flechsig, 1901; Garcia et al., 2018; Grayson & Fair, 2017; Hill et al., 2010; Sydnor et al., 2021; Truzzi & Cusack, 2023). Early maturation in some areas results in limited future plasticity (Hill et al., 2010), which may lead to lower interindividual variability and reduced susceptibility to environmental influences (Gao et al., 2017) and psychopathological factors (Sydnor et al., 2021). We hypothesize that some areas would exhibit early signs of adult-like organization, especially in sensorimotor areas (Gao, Alcauter, Elton, et al., 2015; Sydnor et al., 2021). Additionally, we hypothesize that areas with adult-like organization would overlap with regions showing low interindividual variability in functional network assignments (Dworetsky et al., 2021; Gordon, Laumann, Adeyemo, et al., 2017; Gratton et al., 2018; Hermosillo et al., 2024; Kong et al., 2019; Langs et al., 2016; Seitzman et al., 2019).

In the present work, we used fMRI datasets from two groups: 120 adults (aged 19-32 years) and 181 typically developing infants (aged 8-60 months). We aimed to quantify how well the adult and infant networks describe the modular structure in both adult and infant FC. We also examined if adult networks fit some areas better than others by mapping the spatial distribution of this fit. We identified a subset of areas where infant FC was more similar within their assigned adult networks than within alternative adult networks. We demonstrate the consequences of network model choice by comparing the relationship between chronological age and within-network FC in our area subset to that of all areas. Lastly, we compared the spatial distribution of our area subset to locations with low variance in functional network identity across individuals and demonstrate their proximity to each other. Our findings will help researchers working with infant neuroimaging data understand the pros and cons of using adult and infant functional network models, interpret current results in the literature, and select appropriate models for future research.

## 2. Materials and Methods

### 2.1. Data Collection

#### 2.1.1. Washington University 120 (WU 120)

This dataset has been previously described in detail (Power et al., 2017). Briefly, data were collected from 120 healthy young adult subjects during relaxed eyes–open fixation (60 females, mean age = 25 years, age range = 19–32 years). All subjects were native speakers of English and right-handed. Subjects were recruited from the Washington University community and were screened with a self-report questionnaire to ensure that they had no current or previous history of neurological or psychiatric diagnosis, as well as no head injuries resulting in a loss of consciousness for more than 5 minutes. Informed consent was obtained from all subjects. The study was approved by the Washington University School of Medicine Human Studies Committee and Institutional Review Board.

Structural and functional MRI data were obtained with a Siemens MAGNETOM Trio Tim 3.0-T Scanner (Erlangen, Germany) and a Siemens 12-channel Head Matrix Coil. A T1-weighted sagittal magnetization-prepared rapid acquisition gradient-echo (MP-RAGE) structural image was obtained [time echo (TE) = 3.08 ms, time repetition, TR (partition) = 2.4 s, time to inversion (TI) = 1000 ms, flip angle = 8°, 176 slices with 1 × 1 × 1 mm voxels]. An auto-align pulse sequence protocol provided in the Siemens software was used to align the acquisition slices of the functional scans parallel to the anterior commissure–posterior commissure plane of the MP-RAGE and centered on the brain. This plane is parallel to the slices in the Talairach atlas (Talairach & Tournoux, 1988).

During functional MRI data acquisition, subjects were instructed to relax while fixating on a black crosshair that was presented against a white background. Functional imaging was performed using a BOLD contrast-sensitive gradient-echo echo-planar imaging (EPI) sequence (TE = 27 ms, flip angle = 90°, in-plane resolution = 4 × 4 mm). Whole-brain EPI volumes (MR frames) of 32 contiguous, 4-mm-thick axial slices were obtained every 2.5 s. A T2-weighted turbo spin-echo structural image (TE = 84 ms, TR = 6.8 s, 32 slices with 1 × 1 × 4 mm voxels) in the same anatomical planes as the BOLD images was also obtained to improve alignment to an atlas. Anterior→Posterior (AP) phase encoding was used for fMRI acquisition. The number of volumes collected from subjects ranged from 184 to 724 (mean = 336 frames, 14.0 min).

#### 2.1.2. Baby Connectome Project (BCP)

Full-term (gestational age of 37-42 weeks) infants free of any major pregnancy and delivery complications were recruited as part of the Baby Connectome Project (Howell et al., 2019). All procedures were approved by the University of North Carolina at Chapel Hill and the University of Minnesota Institutional Review Boards. Informed consent was obtained from the parents of all participants. In the final cohort used following fMRI data quality control (described below), we retained 313 fMRI sessions from 181 individuals (95 females, 8-60 months, mean 19.1 months, and standard deviation 8.3 months) during natural sleep (Supplementary Figure 1A). The number of longitudinal points range from 1 to 6: 90 individuals with 1 time point, 60 individuals with 2 time points, 24 individuals with 3 time points, 5 individuals with 4 time points, 1 individual with 5 time points and 1 individual with 6 time points (Supplementary Figure 1B). In a supplementary analysis, we used an additional 15 fMRI sessions collected while participants were watching a movie clip.

All MRI images were acquired on a Siemens 3T Prisma scanner with a 32- channel head coil at the University of Minnesota and the University of North Carolina at Chapel Hill during natural sleep without the use of sedating medications. T1-weighted (TR=2400 ms, TE=2.24 ms, 0.8 mm isotropic; flip angle = 8°), T2-weighted images (TR=3200 ms, TE=564 ms, 0.8 mm isotropic), spin echo field maps (SEFM) (TR=8000 ms, TE=66 ms, 2 mm isotropic, MB=1), and fMRI data (TR=800 ms, TE=37 ms, 2 mm isotropic, MB=8) were collected. A mixture of Anterior→Posterior (AP) and Posterior→Anterior (PA) phase encoding directions was used for fMRI acquisition in each session, but they were concatenated into one time series. A subset of data had a 720-ms TR (N = 95 out of 313 sessions). The number of low-motion volumes collected from subjects ranged from 840 to 2100 (mean = 1306 frames, 16.9 min).

### 2.2. fMRI analysis

#### 2.2.1. MRI data preprocessing

While both datasets were acquired on Siemens 3T scanners, acquisition parameters and preprocessing of the structural and functional MRI data were optimized for each specific cohort. For a detailed comparison of the acquisition, processing, and quality control procedures, please refer to Supplementary Table 1.

##### 2.2.1.1. MRI data preprocessing – WU120

Functional images were first processed to reduce artifacts including (1) Correction of odd versus even slice intensity differences attributable to interleaved acquisition without gaps, (2) correction for head movement within and across runs, and (3) across-run intensity normalization to a whole-brain mode value of 1000. Atlas transformation of the functional data was computed for each individual using the MP- RAGE scan. Each run was then resampled to an isotropic 3-mm atlas space (Talairach & Tournoux, 1988), combining movement correction and atlas transformation in a single cubic spline interpolation (Lancaster et al., 1995).

Additional preprocessing steps were applied to the functional data to reduce the effect of high-motion frames. This was performed in two iterations. In the first iteration, the processing steps were (1) demeaning and detrending, (2), multiple regression including whole-brain, ventricular cerebrospinal fluid (CSF), and white matter signals, and motion regressors derived by Volterra expansion and (3) a band-pass filter (0.009 Hz < f < 0.08 Hz). Following the initial FC preprocessing iteration, temporal masks were created to flag motion-contaminated frames. Motion-contaminated volumes were identified by framewise displacement (FD), defined as the squared sum of the motion vectors (Power et al., 2012). Volumes with FD > 0.2 mm and data segments lasting fewer than 5 contiguous volumes were censored.

The data were then reprocessed in a second iteration, incorporating the temporal masks described above. This reprocessing was identical to the initial processing stream but ignored censored data. Data were interpolated across censored frames using least squares spectral estimation (Power et al., 2014) of the values at censored frames, so that continuous data could be passed through the band-pass filter (0.009 Hz < f < 0.08 Hz) without contaminating frames near high motion frames. Censored frames were ultimately ignored during functional connectivity matrix generation.

Individual surfaces were generated from the structural images and the functional data was sampled to surface space (Glasser et al., 2013). First, following volumetric registration, anatomical surfaces for the left and right hemispheres were generated from each subject’s MP-RAGE image using FreeSurfer’s default recon-all processing pipeline (v5.0)(Fischl, 2012). This pipeline included brain extraction, segmentation, generation of white matter and pial surfaces, inflation of the surfaces to a sphere, and surface shape-based spherical registration of the subject’s “native” surface to the fsaverage surface.

The fsaverage-registered left and right hemisphere surfaces were then brought into register with each other (Van Essen et al., 2012), resampled to a resolution of 164000 vertices using Caret tools (Van Essen et al., 2001) and subsequently down-sampled to a 32492 vertex surface (32k fs_LR). The BOLD volumes were sampled to each subject’s individual “native” midthickness surface (generated as the average of the white and pial surfaces) using the ribbon-constrained sampling procedure available in Connectome Workbench (v0.84) and then deformed and resampled from the individual’s “native” surface to the 32k fs_LR surface. Finally, the time courses were smoothed along the 32k fs_LR surface using a Gaussian smoothing kernel (σ = 2.55 mm).

##### 2.2.1.2. MRI data preprocessing – BCP

MRI data were processed using the DCAN-Labs infant-abcd-bids-pipeline (v0.0.22) largely following steps described previously (Feczko et al., 2021). Structural MRI data underwent HCP-style processing (Feczko et al., 2021; Glasser et al., 2013), including ANTS N4 bias correction, ANTS denoising, T1/T2 distortion correction/registration, and finally ANTS SyN algorithm deformation alignment to an infant MNI template. In addition, a refined brain mask was generated from data that was segmented using in-house age-specific templates via Joint Label Fusion (JLF). The toddler-specific mask and segmentation were substituted into the FreeSurfer (Fischl, 2012) pipeline and used to refine the white matter segmentation and guide the FreeSurfer surface delineation. The native surface data were then deformed to the 32k fs_LR template via a spherical registration.

A scout image (frame 16 in each run) was selected from the fMRI time series for functional MRI preprocessing. The scout was distortion-corrected via spin-echo field maps, served as the reference for motion correction via rigid-body realignment (Feczko et al., 2021), and was registered to the native T1. Across-run intensity normalization to a whole-brain mode value of 10,000 was then performed. These steps were combined in a single resampling with the MNI template transformation from the previous step, such that all fMRI frames were registered to the infant MNI template. Manual inspection of image quality of structural and functional data was conducted to exclude sessions with bad data quality.

To prepare the functional data for FC analysis, further processing steps were applied after sampling the BOLD data to the 32k fs_LR surface space using steps described in 2.2.1.1. First, functional data were demeaned and detrended in time.

Denoising was then performed using a general linear model with regressors including signal and motion variables. Signal regressors included mean CIFTI gray-ordinate time series, Joint Label Fusion (JLF)-defined white matter, and JLF-defined CSF. Motion regressors included volume-based translational and rotational components and their 24- parameter Volterra expansion. The movement of the head was measured by FD and an age-specific respiratory notch filter (0.28-0.48 Hz) was applied to the FD traces and motion parameter estimates to mitigate the effects of factitious head motion due to infant respiration (Fair, 2020; Kaplan et al., 2022). Frames were censored during demeaning/detrending if their post-respiratory filtering FD value exceeded 0.3 mm to generate the denoised beta values in the general linear model. Bandpass filtering was applied using a second-order Butterworth filter (0.008–0.09 Hz). To preserve the temporal sequence and avoid aliasing caused by missing time points during bandpass filtering, interpolation was used to replace missing frames, and residuals were acquired from the denoising general linear model. In addition, zero-padding was applied to both ends of the BOLD data before filtering to minimize the distortions in the edges of the time series. The data were originally minimally spatially smoothed with a geodesic 2D Gaussian kernel (σ = 0.85 mm). A further smoothing with a geodesic 2D Gaussian kernel (σ = 2.40 mm) was applied to give a final effective smoothing of σ = 2.55 mm to match the smoothing used in the adult dataset (WU 120). Finally, the time series were concatenated across all complete and partially completed scan runs with good data quality. The first 7 frames from each run, frames with > 0.2 mm FD post-respiratory filtering (Kaplan et al., 2022), and outlier frames whose across-vertex standard deviation was more than 3 median absolute deviations from the median of the low FD frames were censored and ignored for functional connectivity matrix construction.

#### 2.2.2. Functional Connectivity Matrix Construction

The preprocessed BOLD time series data of each session were parcellated into 333 non-overlapping areas using the Gordon parcellation (Gordon et al., 2016). This choice of parcellation was justified by recent work by our group that demonstrated that the Gordon parcellation had the best fit among a set of adult parcellations and performed comparably to most available infant parcellations in data from infants aged around 8-30 months (Tu et al., 2024). After that, a total number of frames equivalent to 7.2 minutes of data (560 frames for TR = 0.72 and 600 frames for TR = 0.8) were randomly sampled from the full censored time series in each fMRI session. Pearson’s correlation between the parcellated time series was computed to create a 333 x 333 functional connectivity (FC) matrix. This matrix was then Fisher-Z-transformed. The group-average FC matrix was calculated as the mean FC across fMRI sessions.

### 2.3. Infant and Adult Functional Network Schemes

We used the Gordon network assignments (Gordon et al., 2016) for “Adult Networks” (Figure 1A) and Kardan network assignments (Kardan et al., 2022) for “Infant Networks” (Figure 1B). These network assignments were derived from group-average adult and infant FC on the 333 areas using an Infomap community detection algorithm (Rosvall & Bergstrom, 2010) optimized for identifying networks in FC data (Power et al., 2011). Among the 333 areas, some were originally assigned in communities with fewer than 5 areas and considered unassigned (named “None” and “Unspecified”). These areas commonly fall under locations subjected to the biggest susceptibility artifact (Ojemann et al., 1997). We removed them from all analyses and had 286 areas left for the adult networks (“Gordon”) and 328 areas left for the infant networks (“Kardan”).

**Figure 1.**
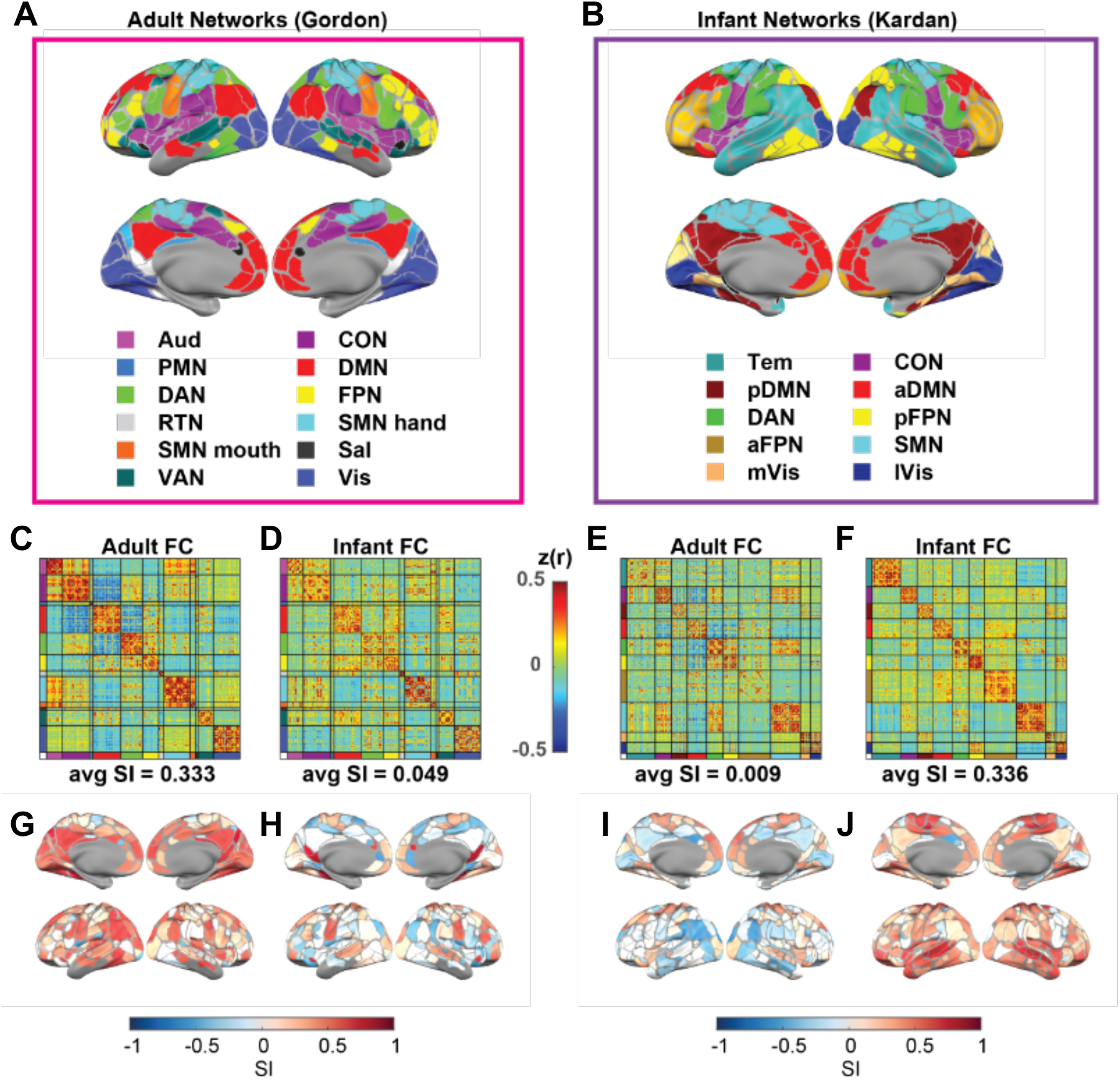
Adult and Infant FC ordered by the adult and infant networks. A) 12 adult networks, B) 10 infant networks, C) average adult FC sorted by adult networks, D) average infant FC sorted by adult networks, E) average adult FC sorted by infant networks, F) average infant FC sorted by infant networks, G) SI of parcels with adult network assignments in adults, H) SI of parcels with adult network assignments in infants, I) SI of parcels with infant network assignments in adults, J) SI of parcels with infant network assignments in infants. Network abbreviations: auditory (Aud), cingulo-opercular (CON), parietal memory (PMN), default mode (DMN), dorsal attention (DAN), fronto-parietal (FPN), retrosplenial temporal (RTN), somatomotor hand (SMN hand), somatomotor mouth (SMN mouth), salience (Sal), and ventral attention (VAN), visual (Vis), somatomotor (SMN), temporal (Tem), posterior frontoparietal (pFPN), posterior default mode (pDMN), lateral visual (lVis), medial visual (mVis), anterior fronto-parietal (aFPN), anterior default mode (aDMN).

The 12 Gordon networks include the auditory (Aud), cingulo-opercular (CON), parietal memory (PMN), default mode (DMN), dorsal attention (DAN), fronto-parietal (FPN), retrosplenial temporal (RTN), somatomotor hand (SMN hand), somatomotor mouth (SMN mouth), salience (Sal), ventral attention (VAN), and visual (Vis) networks. The 10 Kardan networks include somatomotor (SMN), temporal (Tem), posterior frontoparietal (pFPN), posterior default mode (pDMN), lateral visual (lVis), medial visual (mVis), dorsal attention (DAN), anterior fronto-parietal (aFPN), anterior default mode (aDMN).

### 2.4. Functional Network Overlap

The overlap between a network in the Gordon networks and a network in the Kardan networks can be measured with the Dice coefficient, with 0 indicating no overlap and 1 indicating complete overlap. For this analysis, each network is represented with a 333 x 1 vector with 1 for the areas in the network and 0 for the areas outside the network.

### 2.5. Silhouette Index Calculation

Following prior procedures in the literature (Rousseeuw, 1987; Yeo et al., 2011), we calculated the silhouette index (SI) for each area with the correlation distance (i.e. 1- Pearson’s correlation) in the FC profiles:

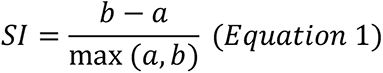

where *b* is the mean between-network correlation distance of the FC profiles, and *a* is the mean within-network correlation distance of the FC profiles. FC profiles here refer to the FC from each area to all other areas (i.e., one row in the FC matrix).

Intuitively, the SI ranges from +1 to -1 with the sign indicating whether the area has a more similar FC profile to areas in its own network (+) or to areas in an alternative network (-). The magnitude indicates the confidence of this assignment, with a higher magnitude suggestive of strong confidence. The average SI for the FC in a network scheme was defined as the average SI across all areas.

We also calculated the silhouette index with the average of all networks rather than just the alternative network in the Supplementary Materials.

To obtain a confidence interval for the average SI across individual sessions, we used a bootstrap 95% confidence interval estimate in 1000 random draws of individual sessions from the full sample (N = 120 for WU120 and N = 313 for BCP) with replacement. The p-value for the average SI different from zero was calculated using the bootstrap distribution of the average SI. The p-value for the difference in the average SI across two samples was calculated using the bootstrapped distribution of the difference in the average SI. Both assumed a two-tailed test.

### 2.6. Identify the Subset of Areas with Similar Network Organization to the Gordon Networks in Infants

A positive SI indicates that the area has a more similar FC profile to areas in its own network. We obtained the subset of areas that had a similar network organization to other areas defined in the Gordon network scheme in infants by only retaining the areas with a positive SI in the group-average infant FC. We refer to this set of positive SI areas as our “area subset”.

### 2.7. Distance Between High Consensus Regions of Interests (ROIs) and the “Gordon Subset” Areas

To quantify the spatial distribution similarity between the locations of low interindividual variability and our area subset, we calculated the Euclidean distance between the high consensus cortical ROIs and the centers of those areas. We used published coordinates of 153 high consensus ROIs calculated previously by identifying locations that demonstrated consistent network assignment across a large majority (i.e. ≥ 75%) of subjects in the Dartmouth dataset (N = 69 subjects, 56 female, average age 20.2 years)(Gordon et al., 2016) when a template-matching procedure (Gordon, Laumann, Adeyemo, et al., 2017) was applied to identify individual network assignments (Dworetsky et al., 2021). The distances in mm were obtained from the ROI locations on a standard adult brain template (a.k.a. MNI coordinates).

For each of the “high consensus” ROIs, we found its distances to the nearest area within and outside our area subset. An average of this difference was recorded and named “distance difference”. A negative distance difference indicates that on average, the high consensus regions were closer to the areas within compared to outside our area subset. To account for the potential effect of differences in the number of areas within (N = 166) and outside (N = 120) our area subset, we repeated the analysis but with the labels of within and outside area subset randomly assigned 1000 times to obtain a null distribution.

### 2.8. Moving Average Analysis Across Age

To examine the FC fit to different network schemes across infancy, we used a moving average analysis across age. For this analysis, we limited our data to the 281 sessions collected at the age of 8-27 months because the data became very sparse and less evenly distributed after 27 months (Supplementary Figure 1A). We first sorted the fMRI sessions by age at scan. Sessions were arranged chronologically by age, and FC averages were computed for consecutive windows of 20 sessions, with each window representing the mean age within it. This window was then shifted by one session at a time until all 281 sessions were accounted for. Subsequently, we calculated the average similarity index (SI) using the same method.

### 2.9. Intraclass Correlation Coefficient

To assess the differences in the reliability of within-network edges for using the Gordon networks with our area subset versus all areas. We quantified the test-retest reliability of FC with the intraclass correlation coefficient (ICC). We assessed the consistency among measurements under the fixed levels of the session factor (Tozzi et al., 2020), referred to as ICC ‘C-1’ (McGraw & Wong, 1996) or ICC (3,1) (Shrout & Fleiss, 1979).

For this analysis, we re-calculated the FC matrices for each individual with two non-overlapping time windows of data from each session. “Test” and “re-test” were defined as the first 6 min and last 6 min of low-motion data, separated by at least 1.2 min low motion data in between to reduce the impact of temporal autocorrelation (i.e. total > 13.2 min low-motion data). Only 167 sessions had enough low-motion data for this analysis. First, the FC values in the upper triangle of each subject’s connectivity matrix were entered as rows in two large matrices (one matrix for “test” and another for “re-test”, one row per subject in each matrix). Then, the corresponding columns of these matrices were compared to obtain an ICC value for each edge. The mean and standard error of the mean of the ICCs within each of the Gordon networks were calculated for our area subset and all areas.

## 3. Results

### 3.1. Adult and Infant functional connectivity clustering was best described by the adult and infant network assignments respectively

Adult networks (”Gordon”) (Figure 1A) and Infant networks (“Kardan”) (Figure 1B) assignments derived using data-driven methods demonstrate a moderate degree of agreement: Normalized Mutual Information (NMI) = 0.5 for the overlapping 281 areas after excluding the “None”/ “Unspecified” network in both adult and infant network assignments. The CON, pDMN, aDMN, SMN, mVis and lVis networks in the Kardan networks tend to have a large dice overlap with a single Gordon network, but Tem, DAN, pFPN, aFPN have a match to multiple Gordon networks (Supplementary Figure 2, network full names defined in section 2.3).

Next, we asked how closely the network assignments matched the similarity of FC profiles within and between different networks and quantified it with the silhouette index (SI; Rousseeuw, 1987; Yeo et al., 2011). We used the average FC across 120 adult sessions and the average FC across 313 infant sessions. We found that adult FC had a more modular organization (Figure 1C) when grouping into adult networks (Figure 1A) than infant networks (Figure 1E). The average SI for areas assigned to adult networks in adult FC (0.333, 95% bootstrap CI = [0.3088, 0.3417], pbootsrap < 0.001) was much higher (pbootsrap < 0.001) than the average SI for areas assigned to infant networks in adult FC (0.009, 95% bootstrap CI = [-0.0015, 0.0168], pbootsrap = 0.042). In contrast, the opposite was observed for infant FC, with a higher average SI (pbootsrap < 0.001) for areas assigned to infant networks in infant FC (0.336, 95% bootstrap CI = [0.3280, 0.3397], pbootsrap < 0.001; Figure 1F) than the average SI for areas assigned to adult networks in infant FC (0.049, 95% bootstrap CI = [0.0406,0.0560], pbootsrap < 0.001; Figure 1D). Furthermore, the results were also qualitatively validated across individual sessions (details in Supplementary Materials), with a much higher SI of adult networks than infant networks on adult FC (Cohen’s *d* = 1.215, p < 0.001) (Supplementary Figure 3C), and a much higher SI of infant networks than adult networks on infant FC (Cohen’s *d* = 2.744, p < 0.001) (Supplementary Figure 4C). Taken together, the adult networks better describe the modular organization in adult FC than infant FC, and the infant networks better describe the modular organization in infant FC than adult FC. However, the SI is comparable for adult FC and infant FC using the best network model, suggesting the presence of modular organization in both cohorts. We also repeated the same analysis for 15 infant sessions scanned while the participants were awake and watching movies and 14 infant sessions scanned during natural sleep with similar age range (Supplementary Figure 5). The difference in average SI between the adult (“Gordon”) networks and infant (“Kardan”) networks on the awake sessions was not significant (pbootsrap = 0.258, paired t-test on individual sessions = 0.71), unlike the difference in asleep sessions with matching age range (pbootsrap <0.001, paired t-test on individual sessions < 0.001). The average SI for areas assigned to adult networks in awake infant FC (0.174, 95% bootstrap CI = [0.1048, 0.1895], pbootsrap <0.001) was much higher (pbootsrap = 0.002) than that in sleeping infant FC (0.084, 95% bootstrap CI = [0.0355, 0.0962], pbootsrap = 0.001), but still lower (pbootsrap < 0.001) than that in adult FC.

Notably, some areas tend to have a positive SI for areas assigned to adult networks regardless of the FC age group (Figure 1G & I). Since the spatial distribution of SI across sessions (Supplementary 3A-B & 4A-B) was relatively consistent, it is unlikely that the low SI magnitude in infants was purely driven by high interindividual variability.

Because the Kardan networks were derived from a subset of BCP sessions used in the present analysis and this might inflate the SI in the infant FC data, we also replicated our analysis with networks derived from an independent 2-year-old dataset with similar acquisition and preprocessing protocols (Tu et al., 2024) (Supplementary Figure 6). The SI of the independently derived infant 2-yearl-old networks was lower than the Kardan networks. This may be partially attributed to the wider age range of the BCP data relative to the toddler data used to derive the 2-year networks. However, it may also suggest that the high average SI with Kardan networks is partially attributable to data leakage. However, importantly, the independently derived infant networks demonstrate significantly higher SI compared to Gordon adult networks when applied to infant FC (pbootsrap < 0.001 for the Tu-12networks and Tu-19networks).

### 3.2. A subset of areas demonstrates adult-like network organization throughout development

An SI above zero for an area indicates that its FC profile is more similar to the FC profiles in areas from the same network than areas from any alternative network within a given network scheme (e.g., adult Gordon networks). Therefore, we selected the subset of areas with an SI above zero when the adult networks were applied to the infant FC (166 in total, Figure 2A, Supplementary Table 2). These areas fell into all 11 out of the 12 Gordon networks (i.e., all except for PMN), with the whole RTN, SMN mouth, and Sal networks retained, and the remaining eight networks partially retained (Figure 2B). We validated that the areas with SI above zero are highly consistent across bootstrap samples, with 156 out of the 166 areas having SI above zero in at least 950 out of 1000 bootstraps (Supplementary Figure 7).

**Figure 2.**
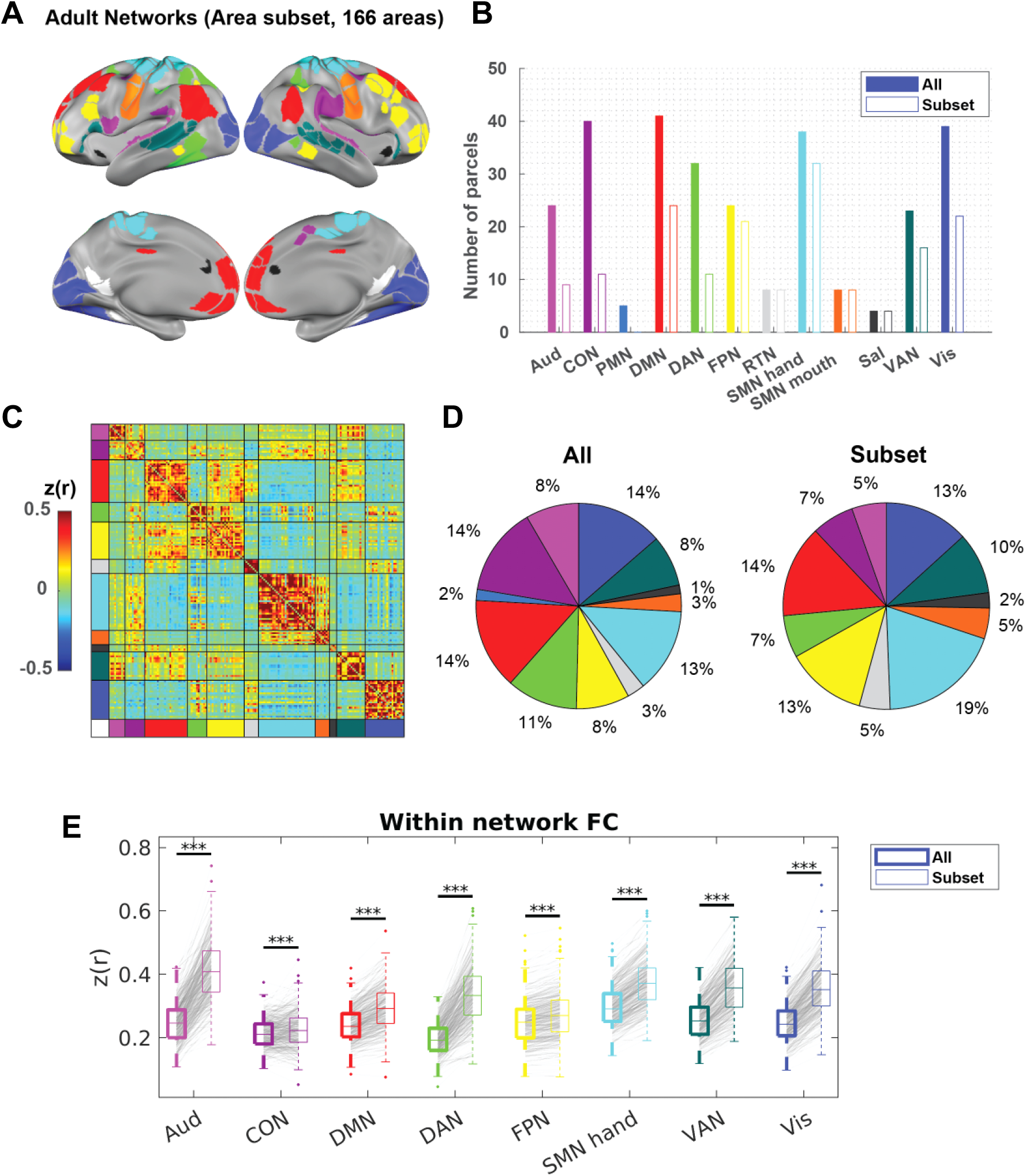
A subset of areas in infants demonstrates adult-like network organization. A) A subset of areas with SI > 0 to adult networks in infants (Figure 1H). B) The number of areas in each adult network with all areas (“All”) or area subset (“Subset”). C) The sorted average FC in infants with our area subset. D) The percentage representation of each network in all areas versus. E) The average within-network FC with all areas (left) versus area subset (right) across individual BCP sessions. FDR-corrected *p* for paired t-tests. *** *p* < 0.001.

As expected, the average SI for areas assigned to adult networks in infant FC was much higher (p_bootstrap_ < 0.001) using our area subset (0.388, 95% bootstrap CI = [0.3792, 0.3925], p_bootstrap_ < 0.001; Figure 2C) than using all areas. In addition, the average SI for areas assigned to adult networks in adult FC was also marginally higher (p_bootstrap_ < 0.001) using our area subset (0.419, 95% bootstrap CI = [0.3925,0.4300], p_bootstrap_ < 0.001) (Supplementary Figure 8A) than using all areas. Our results suggest that this subset captures the part of the adult networks with more within-network consistency in both infants and adults. Compared with all areas, our area subset was disproportionally enriched in the SMN networks (SMN hand and SMN mouth) (Figure 2D). As expected, the within-network FC was significantly higher across infant sessions (paired t-test, FDR-corrected p < 0.05) for all eight partially retained networks when our area subset was used instead of all areas, with little change in variability (Figure 2E).

Similarly, within-network FC was significantly higher across adult sessions (paired t-test, FDR-corrected p < 0.05) for all seven out of eight partially retained networks (Supplementary Figure 8B). Because our data spanned a wide developmental window, we further investigated whether our observation was influenced by chronological age.

We found that within-network FC for all eight partially retained networks was higher in our area subset in most sessions and that there was no correlation (FDR corrected p > 0.05) between the within-network FC differences and age (Supplementary Figure 9). In general, the within-network FC differences between “All” and “Subset” were larger in infants than in adults (Table 1).

**Table 1.**
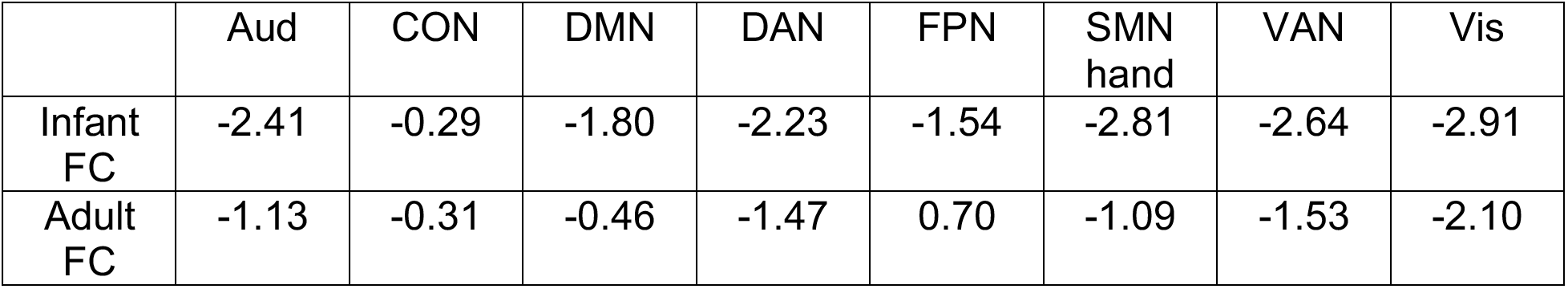
Cohen’s d of the within-network FC differences in All V.S. Subset.

|  | Aud | CON | DMN | DAN | FPN | SMN hand | VAN | Vis |
| --- | --- | --- | --- | --- | --- | --- | --- | --- |
| Infant FC | -2.41 | -0.29 | -1.80 | -2.23 | -1.54 | -2.81 | -2.64 | -2.91 |
| Adult FC | -1.13 | -0.31 | -0.46 | -1.47 | 0.70 | -1.09 | -1.53 | -2.10 |

Specifically, for seven out of the eight partially retained Gordon networks the adult FC was significantly higher (FDR-corrected *p* < 0.05) than the infant FC with all areas (Figure 3A, Table 2). On the other hand, only five out of the eight still demonstrated significantly higher within-network FC when using the subset (FDR-corrected *p* < 0.05) in adults compared to infants, and two out of the eight demonstrated significantly lower within-network FC (FDR-corrected *p* < 0.05) (Figure 3B, Table 2).

**Figure 3.**
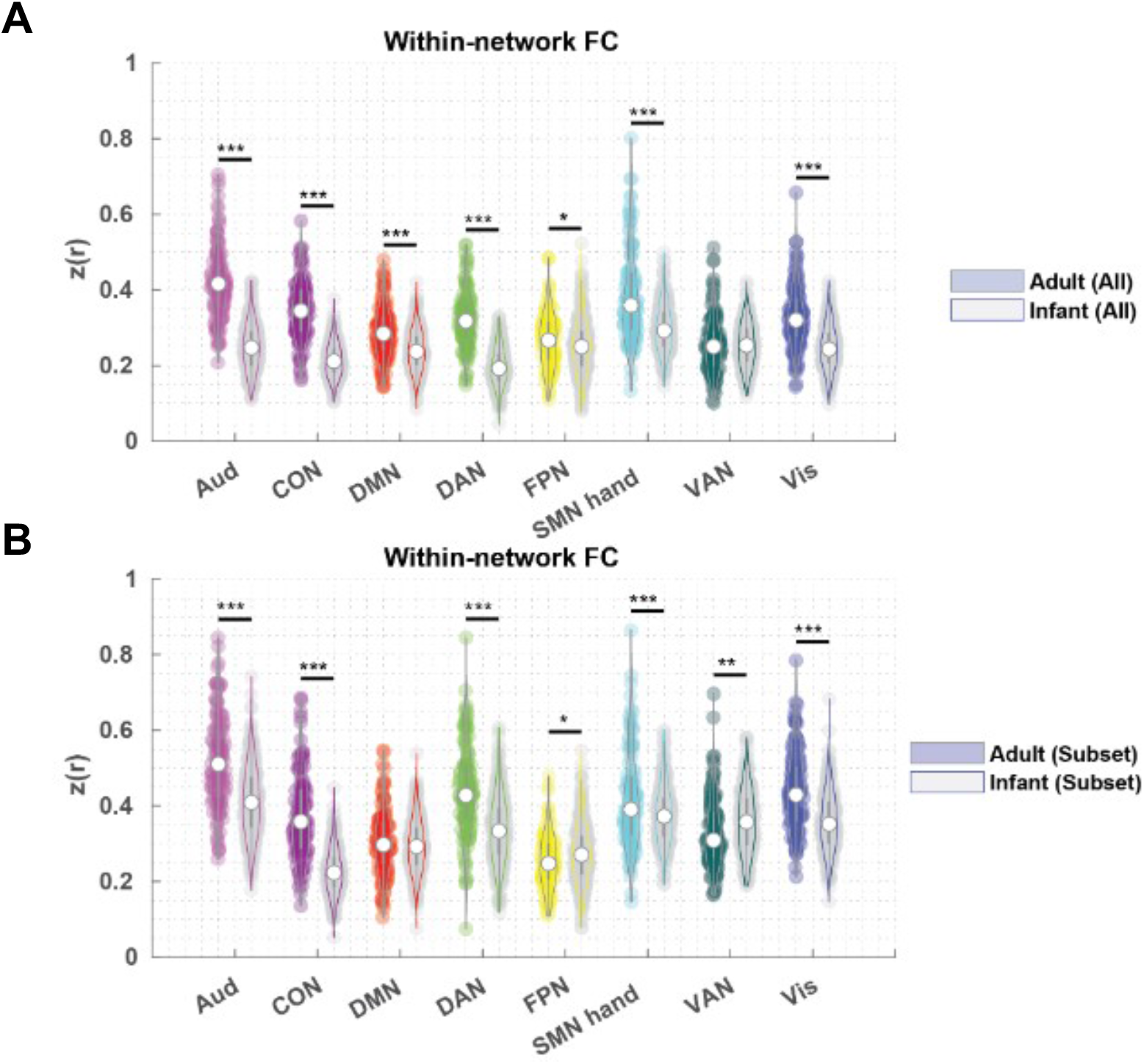
Violin plot of within-network average FC in the Gordon networks. A) within-network average FC using all areas in adults and infants. B) Within-network average FC using our area subset in adults and infants. FDR-corrected *p* for two-sample t-tests. * *p* < 0.05, ** *p* < 0.01, *** *p* < 0.001.

**Table 2.** Cohen’s d of the within-network FC differences in adults V.S. infants.

|  | Aud | CON | DMN | DAN | FPN | SMN hand | VAN | Vis |
| --- | --- | --- | --- | --- | --- | --- | --- | --- |
| All | 2.35 | 2.35 | 0.80 | 2.11 | 0.23 | 1.04 | 0.03 | 1.22 |
| Subset | 1.04 | 1.88 | 0.13 | 1.04 | -0.26 | 0.43 | -0.34 | 0.94 |

Additionally, we found that the effect of chronological age on within-network FC within the infant cohort was also reduced when our area subset was used in place of all areas (Supplementary Figure 10).

Taken together, our results suggest that while the within-network FC within our area subset was higher than that within all areas in both infant and adult datasets, using our area subset would reduce the difference across age.

### 3.3 Adult versus infant FC organizations across 1-2-year-olds

Next, we investigated whether there was any variation between how the different network schemes fit the infant FC at various stages between 1 to 2 years (Gordon adult networks and Kardan infant networks). In addition, we also included a comparison when using our area subset (166 areas, Figure 2A). Using a moving average approach across infant ages, we found a consistent order of the network schemes across 1 to 2 years, with the Gordon (Subset), Kardan (All), and Kardan (Subset) having a similar average SI, and Gordon (All) having a much lower average SI (Figure 4A). Nevertheless, we saw a subtle decrease in difference between adult and infant networks on the infant FC with increasing age. We further investigated this trend using average SI in individual sessions spanning 8 to 60 months (Supplementary Figure 4). We found a weak to moderate negative correlation (r = -0.21, p < 0.001, Supplementary Figure 11A) in infant networks (“Kardan”) fit (average SI) with age, and a weak positive correlation (r = 0.13, p = 0.020, Supplementary Figure 11B) in adult networks (“Gordon”) fit (average SI) with age. Together, this amounted to a moderate negative correlation in the difference in average SI between infant and adult networks with age (Supplementary Figure 11C).

**Figure 4.**
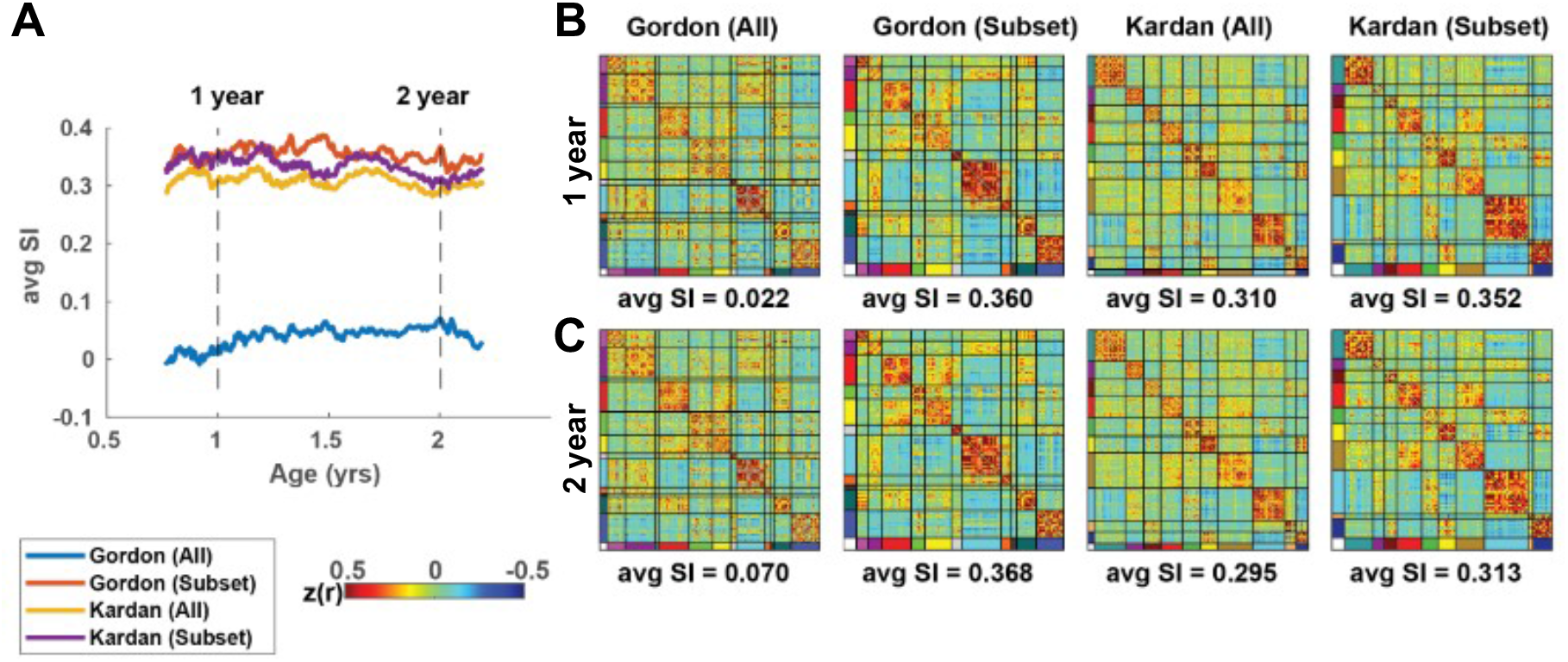
Moving average analysis with the adult networks (“Gordon”) and the infant networks (“Kardan”) using all areas and our area subset. A) Average SI of the average FC in a window for different network assignments. B) Average SI of the window around 1 year old sorted by different network assignments. C) Average SI of the window around 2 years old sorted by different network assignments.

We also replicated Figure 4A with network parcellations derived from an independent dataset to eliminate potential circularity in using the infant networks derived from a subsample of our infant FC data (Supplementary Figure 12).

### 3.4 Our area subset with adult-like network organization is in spatial proximity to the high consensus regions across adult individuals

To quantify the spatial distribution similarity between the locations of low interindividual variability in network identity (“high consensus cortical ROIs”) (Dworetsky et al., 2021) and our area subset, we calculated the Euclidean distance between the centers of the areas within our area subset or alternative areas to the “high consensus cortical ROIs” (Figure 5). Areas within our area subset were 9.5 ± 4.5 mm to the closest high consensus ROIs, whereas areas outside our area subset were 13.5 ± 6.6 mm to the closest high consensus ROIs (Supplementary Figure 13). On average, areas within our area subset were 3.9 mm closer to the “high consensus cortical ROIs”. To rule out the possibility that this difference was driven by the differences in the number of areas within and outside our area subset, we repeated the same analysis by permuting the binary within and outside area subset labels 1000 times to generate a null distribution. We found that the actual difference (3.9 mm) was significantly higher than the null (p < 0.001, permutation testing) (Figure 5C).

**Figure 5.**
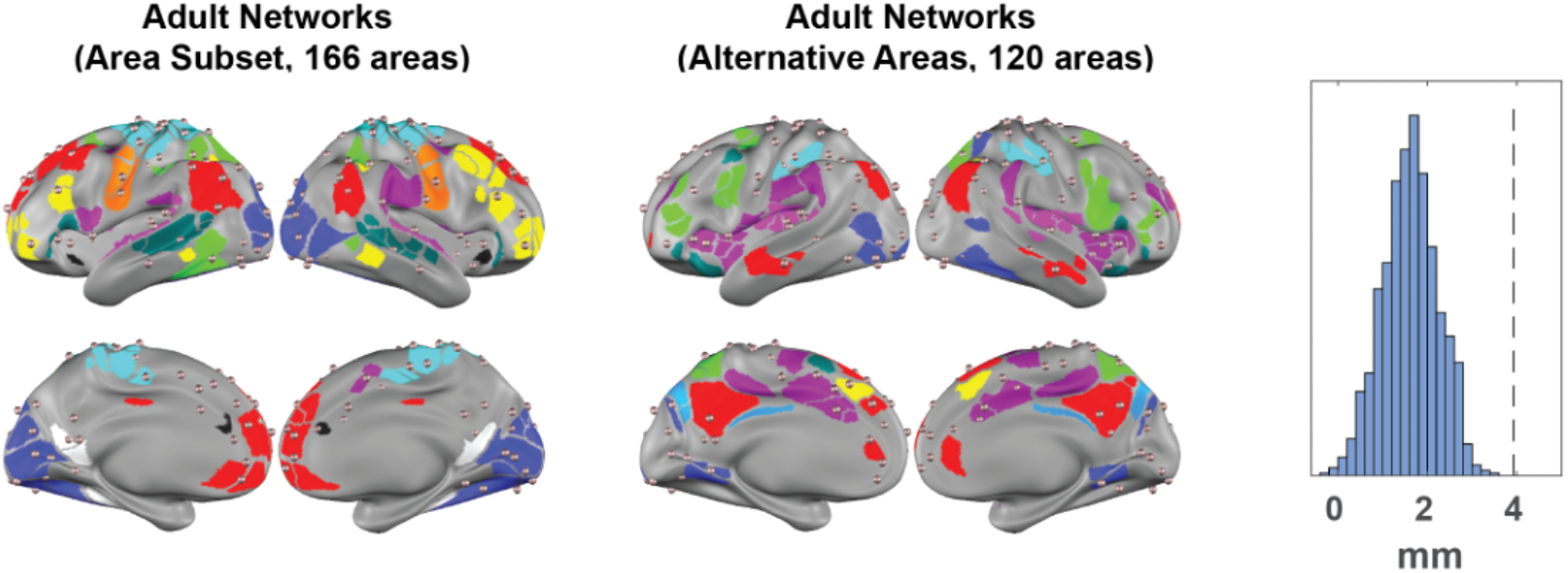
Comparison of area centers to high-consensus regions. A) Our area subset with adult-like network organization (colored surface patches) overlayed with high-consensus regions (rosy brown spheres). B) The areas outside the subset (colored surface patches) overlayed with high-consensus regions (rosy brown spheres). C) The average Euclidean distance between the high-consensus regions and the closest parcel center. Dashed line: the actual difference between the distances in panel A and panel B. Histogram: the difference between the distances with parcels randomly assigned to be in the adult-like (panel A) and not adult-like (panel B) groups 1000 times.

### 3.6. Within-network FC edges in our area subset have a higher test-retest reliability and a higher consistency across subjects

Comparing FC computed from non-overlapping time windows in the same session demonstrated that our area subset had significantly higher within-session reliability than all areas. In particular, four out of eight networks partially retained (Figure 2B) exhibited higher average ICC with our area subset than all areas (two-sample t-test, FDR-corrected *p* < 0.05): Aud (Cohen’s *d* = 0.68), DMN (Cohen’s *d* = 0.13), DAN (Cohen’s *d* = 0.55), and VAN (Cohen’s *d* = 0.43) (Figure 6). To examine whether the contribution of FC edges to individual identification varied across the within- and between-network blocks by the three network schemes, we also quantified the FC group consistency (*ϕ*) and differential power (DP) (Finn et al., 2015). We found the FC- edges connecting our area subset tend to be the ones more consistent across scans and subjects (Supplementary Materials S4, Supplementary Tables 3-6, Supplementary Figure 14).

**Figure 6.**
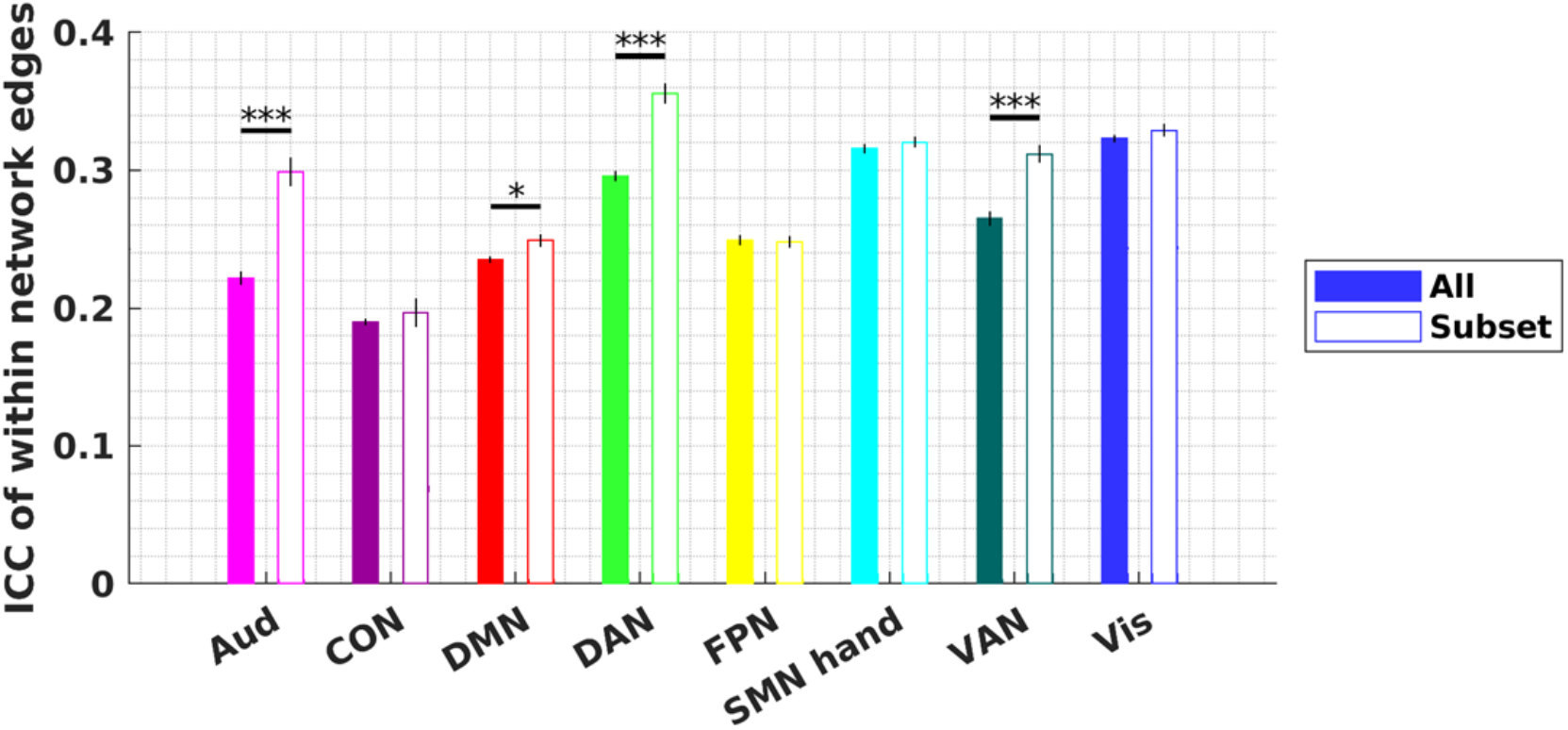
Reliability of within-network edges using our area subset versus all areas. Reliability of within-network edges across two non-overlapping 6-minute windows. Bars show the mean of the ICC and error bars show the standard error of the mean. FDR- corrected *p* for two-sample t-tests. * *p* < 0.05, ** *p* < 0.01, *** *p* < 0.001.

## 4. Discussion

### 4.1. Similarity and differences between adult and infant network organization

We observed that for infants at 8-60 months, adult networks captured some of the features in the infant FC (average SI>0), although much weaker than that in adult FC. Furthermore, it seemed that instead of being less modular and more random, the infant FC data were better described with notably different but related modular organization, including fragmented anterior and posterior segments of higher-order association networks (Eggebrecht et al., 2017; Eyre et al., 2021; Kardan et al., 2022; Marrus et al., 2018).

### 4.2. Identification of a subset of areas that are stable across development

We found that a subset of the areas tended to exhibit more of an adult-like network FC clustering pattern even in infants, forming the cores of adult networks. While substantial attention has been devoted to the differences in infant network organization throughout development (F. Wang et al., 2023; Wen et al., 2019, 2020), it is also desirable to note their similarities to older children and adults (Fransson et al., 2007; Gao, Alcauter, Elton, et al., 2015). Here we observed that the difference in within-network FC across ages was reduced when an area subset was used instead of all areas. The FC within our area subset was also more consistent both within the session and across sessions. These areas likely form the early scaffold for what will eventually become the adult networks (Grayson & Fair, 2017).

### 4.3. The possible role of childhood experience in shaping the interindividual differences in functional networks

Our results suggest that interindividual variability in functional network topography might have a developmental origin. To a first approximation, the spatial topography of our area subset closely matches the locations with low interindividual variability in network identity (Dworetsky et al., 2021; Gordon, Laumann, Gilmore, et al., 2017; Hermosillo et al., 2024). On the other hand, the other areas approximately correspond to previously identified hub nodes (Power et al., 2013) and integration zones (Hermosillo et al., 2024). One potential explanation for this observation is that some parts of the brain mature earlier than others in terms of functional network organization. There might be biological or evolutionary reasons that parts of the adult networks mature later, such as to devote prenatal resources to regions most important for early survival (Hill et al., 2010). In addition, the regional variability of network stability might also be linked to variability in the expression of excitatory and inhibitory features across the cortex (Sydnor et al., 2021). The idea that areas with higher FC variability have more significance in interindividual differences is further reinforced by research demonstrating that FC in cortical areas with high FC variability has more predictive power in behavioral and cognitive domain features (Mueller et al., 2013). It is important that future researchers recognize the regional variability in functional network stability. This variability may serve as a useful biomarker for psychopathology (Sydnor et al., 2021), or may guide targeted brain stimulation interventions.

We did not observe a strong bias in the over-representation of sensorimotor networks compared to association networks in our area subset, despite the literature suggesting that sensorimotor networks mature earlier than association networks (Gao, Alcauter, Elton, et al., 2015; Sydnor et al., 2021). Our area subset spanned both sensorimotor and association networks along the functional hierarchy of the neocortex (Flechsig, 1901; Mesulam, 1998; Sydnor et al., 2021). One potential limitation is that our infant cohort was older than eight months and significant earlier neurodevelopmental changes along the sensorimotor-association hierarchy might have happened before eight months (Bethlehem et al., 2022; Flechsig, 1901). Another possibility is that the sensorimotor functional networks definition was inaccurate, e.g. the auditory network might have incorporated parts of secondary somatosensory regions (Raju & Tadi, 2024), leading to heterogeneous FC profiles within the auditory network

### 4.4. Using an area subset to improve statistical power and interpretability

There are pros and cons of using a pre-existing functional network model and a data-driven functional network model for infant neuroimaging research. Studies of functional networks in infants have often implemented unsupervised methods (i.e. clustering or similar types of community detection algorithms) to find age-specific modules and called them “functional networks” (Eggebrecht et al., 2017; Kardan et al., 2022; Marrus et al., 2018; Molloy & Saygin, 2022; Myers et al., 2024; Sylvester et al., 2022; F. Wang et al., 2023; Wen et al., 2019, 2020). These identified modules were by definition a good representation of the organizational structure in the data and may help alleviate the problem of reproducibility in brain-wise association studies (Hermosillo et al., 2024; Marek et al., 2022). However, unlike the adult networks that have been extensively validated with task fMRI data to corroborate their “functional” roles (Power et al., 2011; Wig, 2017; Yeo et al., 2011), those age-specific modules often lack biological support for their functions, reducing their relevance to a broader developmental context. On the other hand, using the adult-network topography directly on infants neglects the infant-specific organizational features and risks including spurious variability in neuroimaging measurements (be they from fMRI, PET, or fNIRS), leading to reduced effect size and statistical power (Hermosillo et al., 2024), or an exaggerated difference across development. For example, our results in section 3.2 suggest that differences in within-network FC across age groups might be partially attributed to the misspecification of functional network identity.

Here, we propose an alternative strategy that uses an area subset that is relatively stable throughout development. This approach strikes a balance between interpretability/comparability across cohorts and reliability/reproducibility. This idea of using a subset of the brain areas to define ROIs as an approach to improve statistical power has been mentioned in prior literature (Dworetsky et al., 2021; Hermosillo et al., 2024). However, instead of focusing on the subset of brain areas with interindividual variability, we focus on excluding the subset of brain areas that had a potential network misidentification in the infant cohort. Alternatively, one might be interested in focusing on the areas that are unstable across development depending on the research question at hand, which may have behavioral or clinical significance.

### 4.5. Implications for Precision Functional Mapping in Developmental Cohorts Using Adult Group Priors

As demonstrated in our results, on average the adult functional networks did not well represent the organization of infant FC into internally similar clusters. This observation might have important implications for research using an adult functional network model to generate individual-specific functional networks in pediatric cohorts (Hermosillo et al., 2024; Moore et al., 2024; Sun et al., 2023). Motivated by recent research findings showing that functional network topography across human individuals qualitatively differs from group-average estimates (Gordon, Laumann, Gilmore, et al., 2017; Gratton et al., 2018; Laumann et al., 2015), researchers have emphasized the importance of precision functional mapping (Gratton et al., 2022). However, reliable identification of individualized functional networks with unsupervised clustering or community detection procedures requires extended data acquisition. For example, with the Infomap algorithm (Power et al., 2011; Rosvall & Bergstrom, 2008), more than 90 minutes of data are required to achieve an average network overlap dice coefficient of > 0.75 (Gordon, Laumann, Gilmore, et al., 2017). Therefore, researchers have developed several semi-supervised methods using adult networks as priors to derive individual functional networks in data with shorter acquisition time (Cui et al., 2020; Gordon, Laumann, Adeyemo, et al., 2017; Hacker et al., 2013; Kong et al., 2019; D. Wang et al., 2015). However, these approaches generally assume that the individual functional networks are highly similar to the adult group average template. This assumption might not be suitable for developmental cohorts. As we demonstrate in this paper, the adult functional networks poorly represent the organization of the infant FC into internally coherent clusters. Two unwanted consequences might arise from this observation. First, the network templates generated by averaging the FC profiles within a poorly defined network might be noisy and inaccurate. Second, the algorithms may incorrectly force a categorical label for locations that poorly match all available networks. Future studies using adult-based priors in developmental cohorts should keep those limitations in mind and develop strategies to mitigate them.

### 4.6. Limitations and Future Directions

The differences in state (asleep in infants and awake in adults) may contribute to the worse fit of adult networks to infant FC. Sleep and the level of arousal are known to modify the FC structure in adults (Chang et al., 2016; Mitra et al., 2017; Tagliazucchi et al., 2012), and FC patterns in sleeping 6- and 12-month-old infants more closely resemble FC patterns in sleeping adults (Mitra et al., 2017). Based on our results from a small sample of awake fMRI in 36-to-60-month-olds, the difference in the quality of clustering between awake adult and awake infant FC using the adult Gordon networks is smaller than the difference in sleeping infant FC. Taken together with the existing literature, these results tentatively support the hypothesis that infant network clustering quality is driven by the interaction of both brain development and sleeping state.

Other differences in the acquisition and processing of the two datasets might introduce further confounds. Additionally, while we observed minimal effects of age on within-network FC, this could be due to the narrow age range of our sample (mostly between 1 and 3 years). The mixing of fMRI signals within the ill-defined areas using an adult parcellation might contribute to the low SI observed in some of those areas which could be improved with the use of infant specific areas (Tu et al., 2024).

Future studies could examine the cellular, molecular, and genetic properties of the areas that have already developed an adult-like organization in infancy to fully understand the biological underpinning of our observation. Furthermore, future studies with larger samples and well-defined behavior measures can explicitly test our deduction that the use of our area subset could improve statistical power and reproducibility for brain-wide association studies. Moreover, it is worth investigating whether the topography and diversity of thalamocortical projections would relate to the variability of functional network stability across the neocortex.

### 4.7. Conclusions

We found that there exists a subset of cortical areas whose FC profiles demonstrate adult-like network organization even in infants, despite the noticeable differences in FC organization between infants and adults. These areas were spatially closer than alternative areas to previously described locations of high network identity consensus in adults. Additionally, within-network FC defined with our area subset was higher in magnitude and more reliable across scans, individuals, and chronological age. We propose the use of adult networks defined by our area subset as a complementary approach of studying infant FC than using age-specific functional networks derived from data-driven methods. This would strengthen reliability, yet at the same time encourage interpretability and comparability across developmental stages. The biological basis of our observations as well as their psychopathological and behavioral impacts may become interesting topics for future research.

## Author Contributions

JCT conceptualized the project. JCT and YW conducted a formal analysis. OK, OM, TKMD, CMS, DD, XW, and YW processed/curated the data. MDW, JCT, JTE were responsible for funding acquisition. JCT and MDW wrote the original draft. Everyone contributed to the review and editing of the final manuscript.

## Data and Code Availability

The WU 120 data can be downloaded from https://legacy.openfmri.org/dataset/ds000243/.

The BCP data can be downloaded from the National Institute of Mental Health Data Archive (NDA) at https://nda.nih.gov/edit_collection.html?id=2848. The preprocessing scripts are available at https://github.com/DCAN-Labs/dcan-infant-pipeline. The analysis scripts used to generate the results and figures are available on https://github.com/cindyhfls/Tu-2024-GordonSubset-DCN.

The test for the difference between correlations were implemented from the function corr_rtest downloaded from MATLAB central. The Intraclass Correlation Coefficient (ICC) was calculated from the function ICC downloaded from MATLAB central.

## Acknowledgement

The authors would like to thank Ari Segel for assistance in statistical analysis. The authors would also like to thank all families participated and technicians who have helped with the collection, processing, and curation of the data.

## Declaration of generative AI and AI-assisted technologies in the writing process

During the preparation of this work the authors used ChatGPT in order to improve sentence structure and language precision. After using this tool/service, the authors reviewed and edited the content as needed and take full responsibility for the content of the publication.

## Supplementary Materials

### Supplementary Results

#### S1. Silhouette index in individual sessions

We calculated the silhouette index in group-average data because individual sessions are noisy and have a relatively short acquisition time. However, our group level results were consistent with results obtained by calculating silhouette index in individual sessions (Supplementary Figures 3-4). Specifically, the average SI (−0.0054 ± 0.0714) was not different from zero in individual adult sessions (two-tailed t-test, p = 0.41) for adult (“Gordon”) networks. On the other hand, the average SI (−0.0859 ± 0.0318) was significantly smaller than zero in individual adult sessions for infant (“Kardan”) networks (two-tailed t-test, p<0.001). In addition, the average SI (−0.0636 ± 0.0492) was smaller than zero in individual infant sessions (two-tailed t-test, p < 0.001) for adult(“Gordon”) networks. On the other hand, the average SI (0.0492 ± 0.0472) was significantly greater than zero in individual adult sessions for infant (“Kardan”) networks (two-tailed t-test, p<0.001).

#### S2. Silhouette index of adult networks in infant FC with the mean in all alternative networks

By default, the silhouette index compares the current network to the best alternative network, which also depends on the quality of alternatives. However, other researchers have chosen to use a similar metric that compares the average within-network similarity to the average between-network similarity across all alternative networks, rather than just the best alternative (Ji et al., 2019). This approach tends to be less conservative and generally results in a higher silhouette index when calculated in this manner.

When the SI was calculated using the mean in all alternative networks rather than the mean of the best alternative network, they were still moderately correlated with the SI reported in the main results (Pearson’s r = 0.74, *p* <0.001). However, since the mean of similarity to all alternative networks (especially to the ones spatially distant from the area in question) would tend to be lower than the best alternative, the SI is positively shifted with almost all parcels having SI > 0 (Supplementary Figure 15).

#### S3. Age effect on within-network (Gordon networks) FC is smaller in magnitude with our area subset than with all areas

To test the hypothesis that our area subset has relatively stable within-network FC across chronological age in infants, we compared the age effect on within-network FC when the networks include only our area subset versus all areas. The age effect of within-network FC was quantified with a Spearman’s correlation (ρ). The significance of the difference between the correlation between chronological age and within-network FC in our area subset versus all areas is calculated with a Z-test on Fisher-Z-transformed r values.

We additionally examined the within-network FC in infants across chronological age. We computed within-network FC across age using full versus our area subset. For the eight networks that were partially retained, five networks demonstrated a significant correlation between within-network FC and age (*p* < 0.05, Spearman’s *ρ*): the within-network Aud, SMN hand and Vis networks were negatively correlated with age and the within-network FC in DAN and the FPN were positively correlated with age. The age effect was greater in magnitude with the full set of areas (Figure 3A) than with only the partially retained areas (Figure 3B) for the SMN hand network, although not significant when comparing the Fisher-Z-transformed *ρ* values (*Z* = 1.588, one-sided p = 0.056). Similar results were found for other networks, where the age effect was less negative for Aud, SMN hand and Vis networks, and less positive for DAN and FPN, but none of them had a significant (p < 0.05) Z-test. To examine the robustness of our result to the selection of data samples, we generated 1000 bootstrapped samples of the infant sessions. We found that the sign of the difference was consistent across bootstrap samples (i.e., on average the networks using our area subset was less correlated with age than all areas) (Figure 3C). The mean and 95% confidence interval for the bootstrap showed a mean difference in Fisher-Z-transformed *ρ* values for full versus subset was -0.1139 [-0.1721,0.0129] for Aud, -0.1386 [-0.1684, -0.0814] for SMN hand, 0.0020 [-0.0887, 0.0348] for Vis, -0.0205 [-0.0120, 0.1439] for DAN and 0.0089 [0.0071, 0.0511] for FPN (Figure 3C).

#### S4. Group consistency and differential power of FC edges

Prior studies suggested that it was possible to identify individuals using FC in infants from the BCP dataset (Hu et al., 2022; Kardan et al., 2022). To assess which FC edges (i.e. connections between a pair of areas) are more consistent across individuals versus distinct across individuals, we calculated the group consistency (*ϕ*) and differential power (DP) measures (Finn et al., 2015). We aim to describe the distribution of highly consistent edges and highly differentiating edges with respect to adult and infant network models. For this analysis, we only use the one session from each of the 115 unique subjects with at least 13.2 min low-motion data. Given two sets of connectivity 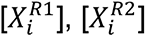 obtained from the two resting scan windows (*R1* and *R2*) after z-score normalization, the edgewise product vector φ*_i_* was computed as

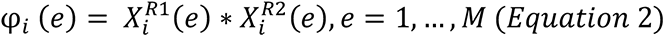

where *i* indexed the subject, *e* indexed the edge, and *M* indexed the total number of FC edges. The sum of φ*_i_* over all edges is the correlation between 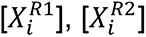. The group consistency *ϕ* was computed as the mean of φ*_i_* across all subjects. We defined the edges with the top 10% *ϕ* values to be “highly consistent”.

Similarly, the edgewise product vector φ*_ij_* was calculated between patterns from different subjects, for example:

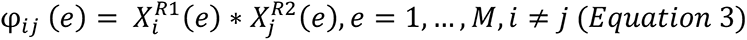

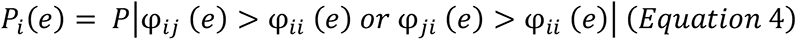

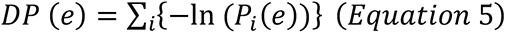

We defined the edges with the top 10% DP values as “highly differentiating”.

Consistent with the findings in previous literature, we observed that a large percentage (∼50%) of FC edges in the within-network blocks tend to be highly consistent. On the other hand, much fewer FC edges in between-network blocks (∼6%) were highly consistent (Supplementary Figure 14; Supplementary Table 3). The sensorimotor networks especially had a large proportion of highly consistent within-network FC edges (Supplementary Table 4). Moreover, using adult networks defined by our area subset, the percentage of highly consistent edges within networks increased substantially for all eight partially retained networks (Supplementary Table 4), indicating that the adult network spanned by our area subset over-represented areas with highly consistent FC between them.

On the other hand, within-network blocks tend to have only a slightly larger percentage of highly differentiating FC edges (∼15%) than between-network blocks (∼10%) (Supplementary Table 5-6), with both increased and decreased proportion of highly differentiating edges when using our area subset instead of all areas.

---

**Supplementary Figure 1.**
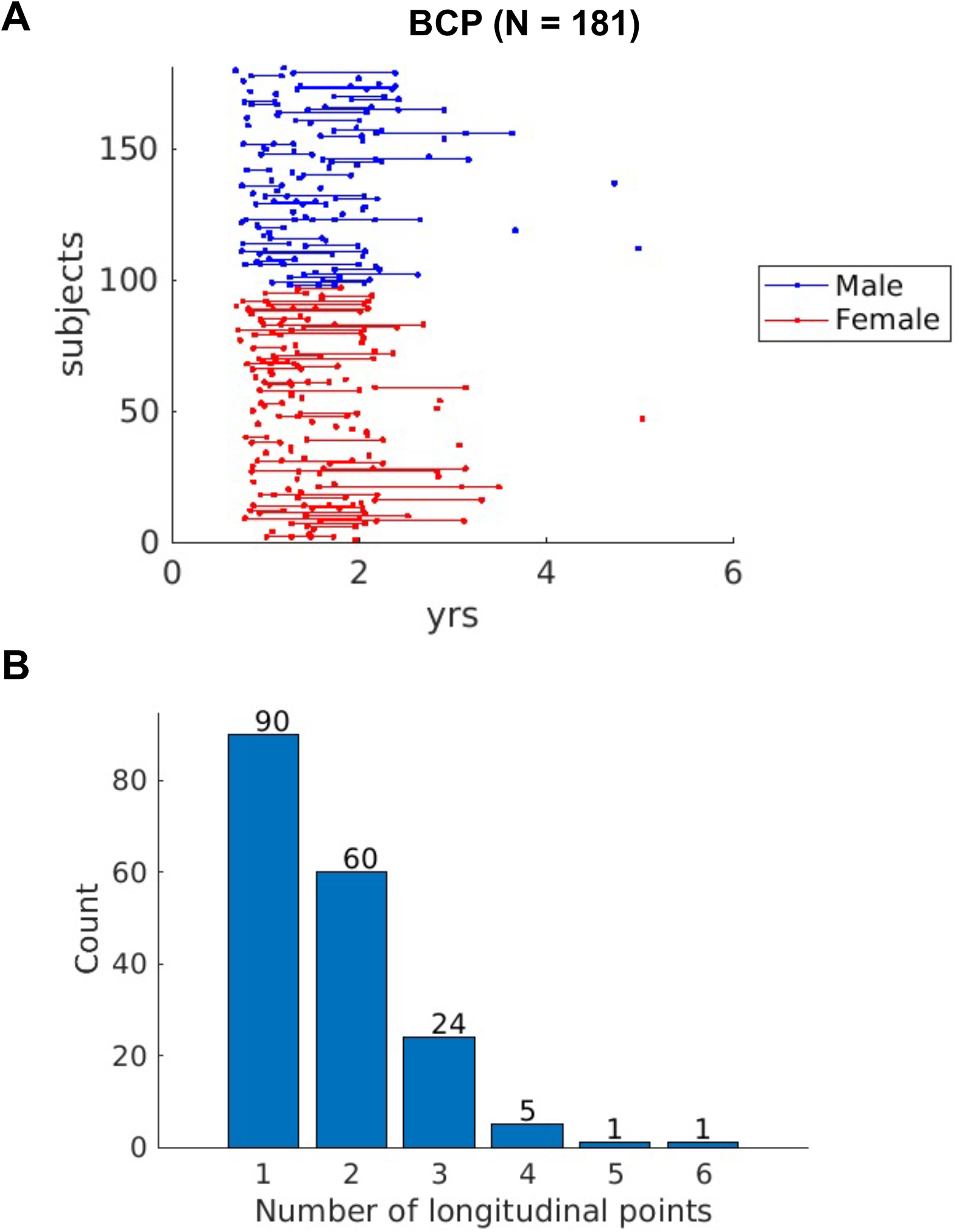
Distribution of age and sex of individual infants in the BCP dataset. A) The age time points of 181 infants ordered by sex. B) The count of number of individuals with 1-6 longitudinal points.

**Supplementary Figure 2.**
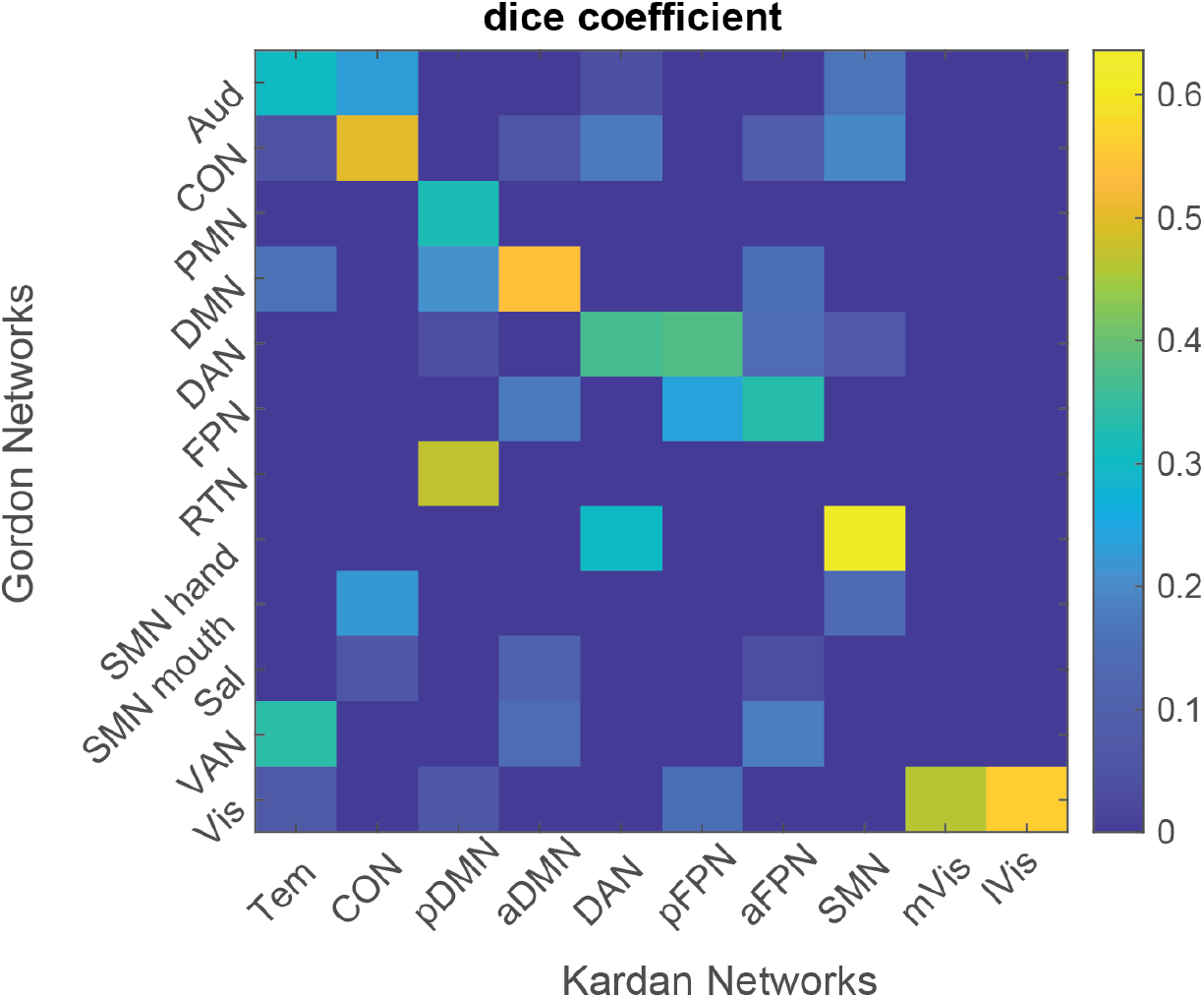
Dice overlap between Adult Networks (“Gordon”) and Infant Networks (“Kardan”). Network abbreviations: auditory (Aud), cingulo-opercular (CON), parietal memory (PMN), default mode (DMN), dorsal attention (DAN), fronto-parietal (FPN), retrosplenial temporal (RTN), somatomotor hand (SMN hand), somatomotor mouth (SMN mouth), salience (Sal), and ventral attention (VAN), visual (Vis), somatomotor (SMN), temporal (Tem), posterior frontoparietal (pFPN), posterior default mode (pDMN), lateral visual (lVis), medial visual (mVis), anterior fronto- parietal (aFPN), anterior default mode (aDMN).

**Supplementary Figure 3.**
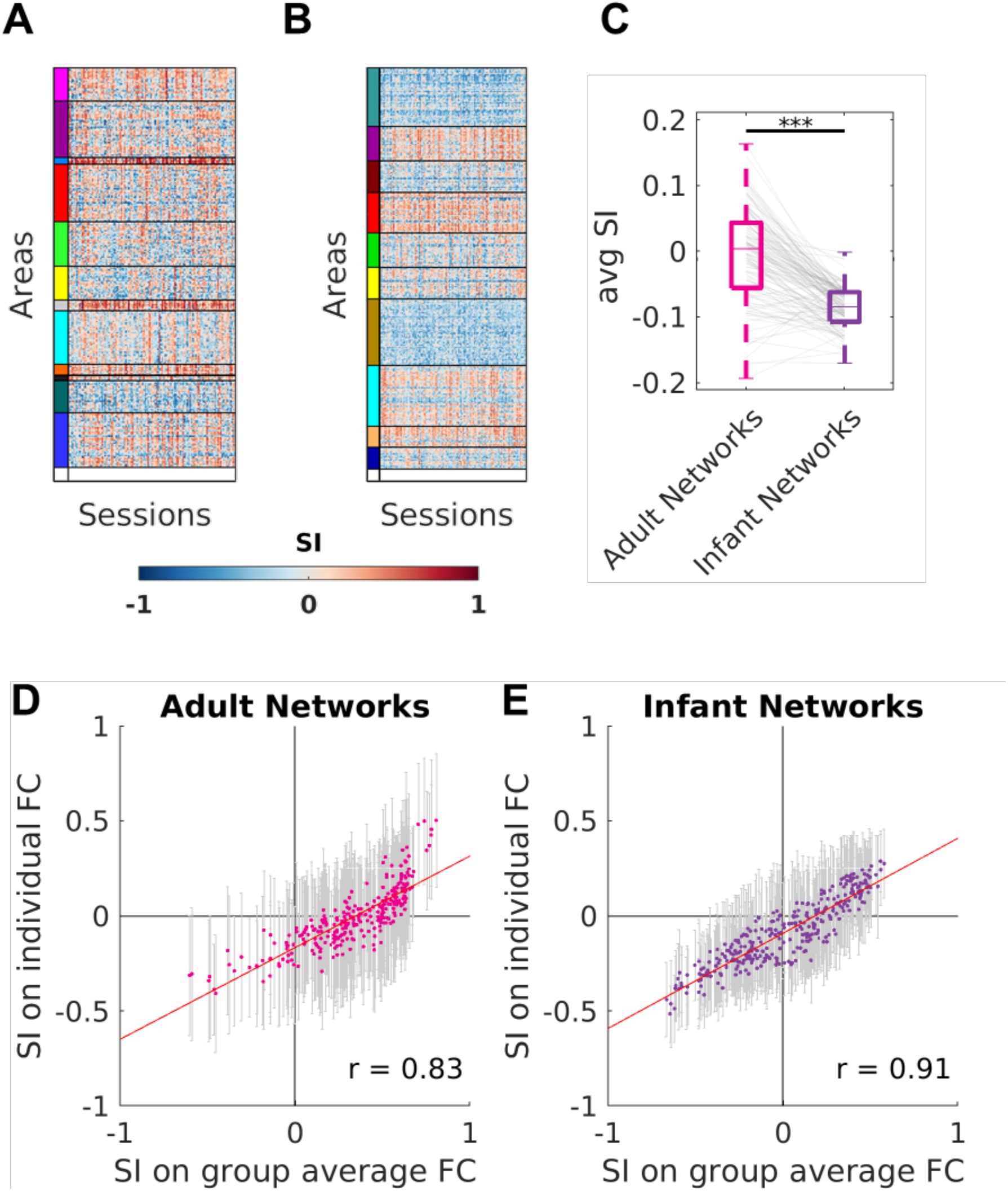
Silhouette index (SI) of adult and infant networks on individual adults’ FC. A) SI across adult networks (“Gordon”, 286 areas). B) SI across infant networks (“Kardan”, 328 areas). C) average SI of adult and infant networks across areas on individual adults’ FC. *** p < 0.001 in paired t-test. D) Pearson’s correlation of SI of adult networks on group average FC and the mean of SI on individual FC across 286 areas. E) Pearson’s correlation of SI of infant networks on group average FC and the mean of SI on individual FC across 328 areas.

**Supplementary Figure 4.**
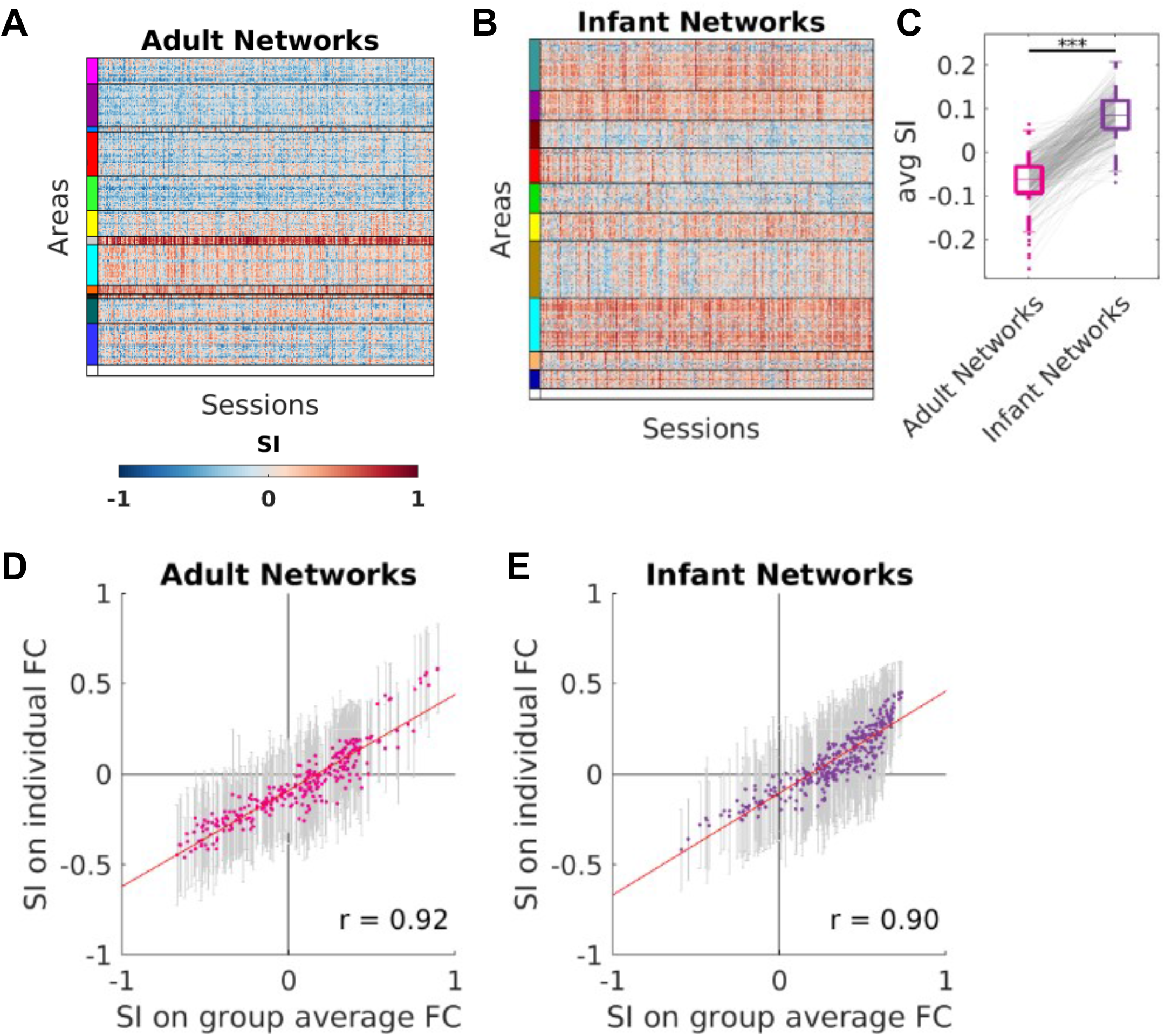
Silhouette index (SI) of adult and infant networks on individual infants’ FC. A) SI across adult networks (“Gordon”, 286 areas). B) SI across infant networks (“Kardan”, 328 areas). C) average SI of adult and infant networks across areas on individual infants’ FC. *** *p* < 0.001 in paired t-test. D) Pearson’s correlation of SI of adult networks on group average FC and the mean of SI on individual FC across 286 areas. E) Pearson’s correlation of SI of infant networks on group average FC and the mean of SI on individual FC across 328 areas. Sessions in A and B are sorted by increasing age from left to right.

**Supplementary Figure 5.**
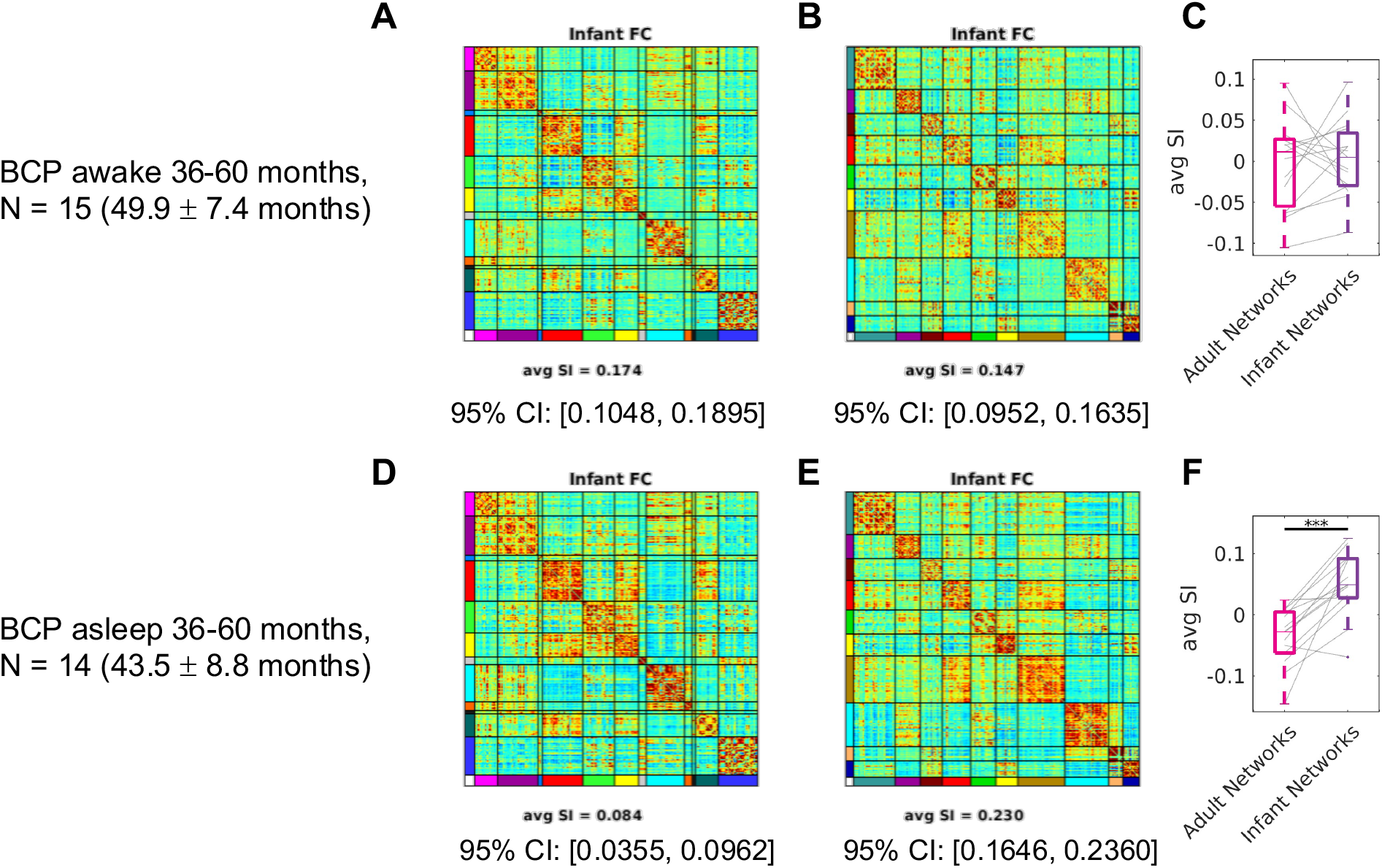
Awake V.S. sleeping infant FC organized by adult (“Gordon”) and infant (“Kardan”) networks. A) The average FC for 15 BCP sessions sorted by adult networks (“Gordon”). B) The average FC for 15 BCP sessions sorted by infant networks (“Kardan”). C) Average silhouette index for individual sessions. D-F) Same as A-C, but for 14 BCP sessions also in approximately the same age range (36-60 months). *** *p* < 0.001 in paired t-test.

**Supplementary Figure 6.**
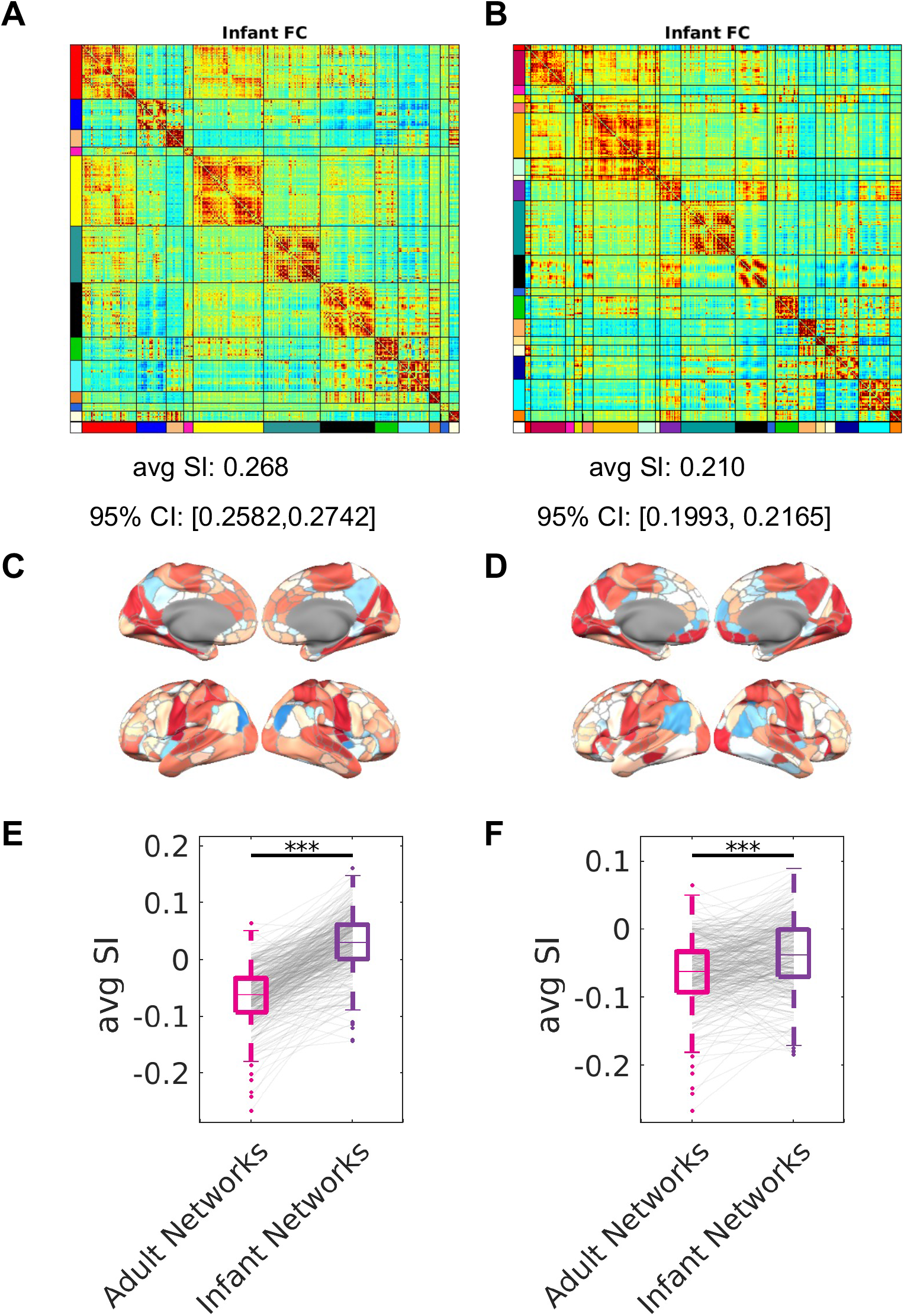
Infant FC sorted by the Tu (326) 12 and 19 networks. A) The average FC for 313 BCP sessions sorted by Tu (326) 12 networks. B) The average FC for 313 BCP sessions sorted by Tu (326) 19 networks. C-D) Silhouette index for each area parcel for A-B. E-F) Average silhouette index for individual sessions. *** *p* < 0.001 in paired t- test.

**Supplementary Figure 7.**
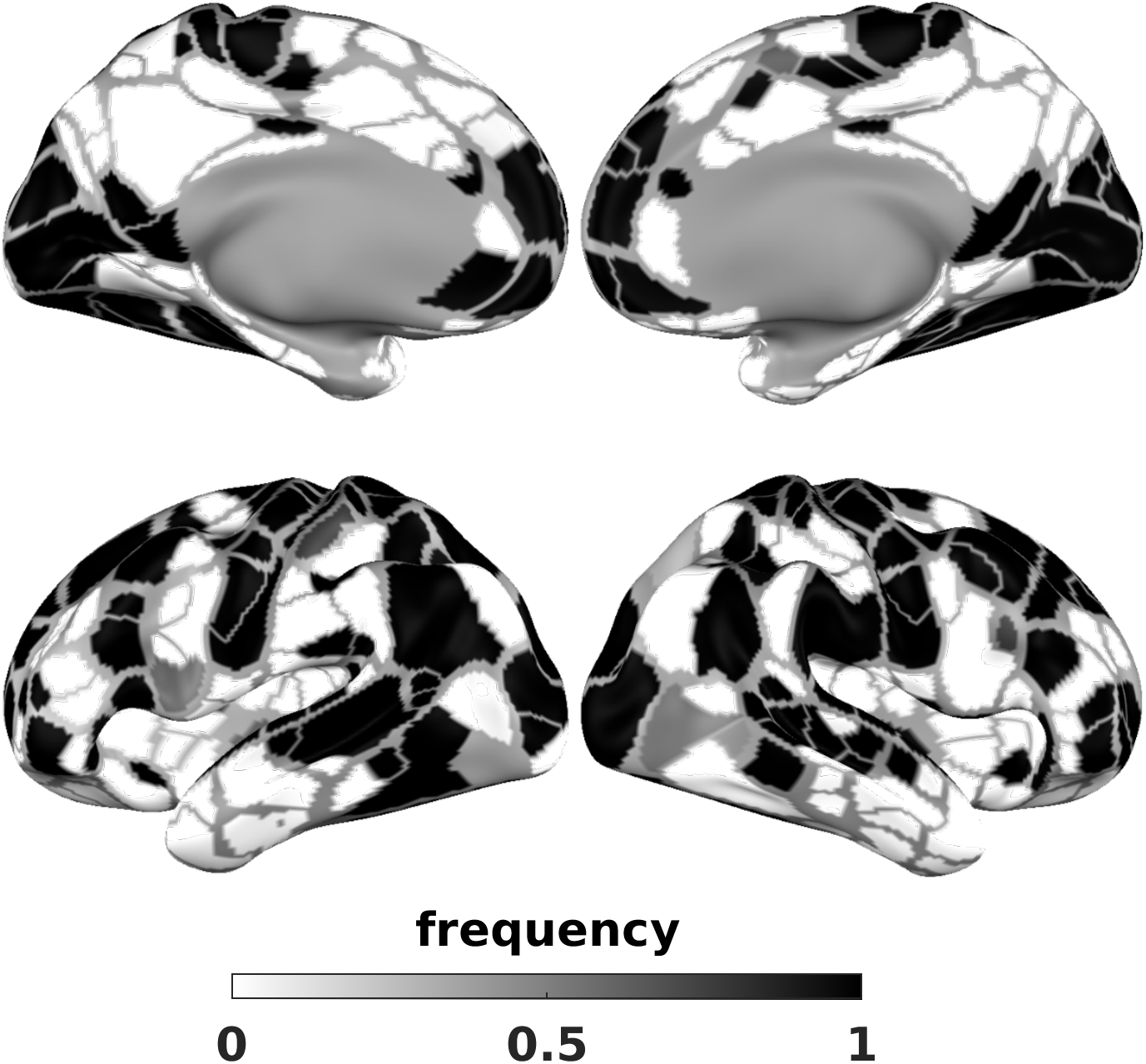
Frequency of SI > 0 across 1000 bootstraps.

**Supplementary Figure 8.**
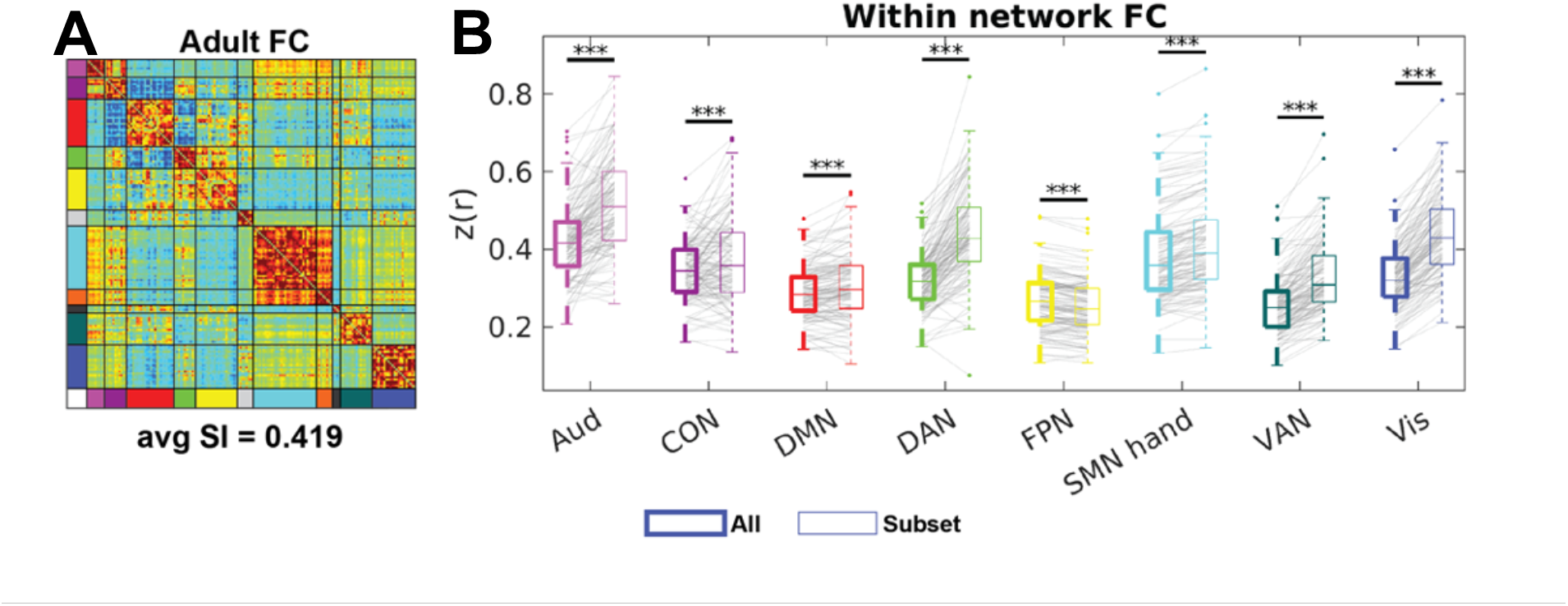
Adult FC using our area subset. A) The sorted average FC in adults with our area subset. B) The average within-network FC with all areas (left) versus our area subset subset (right) across sessions.

**Supplementary Figure 9.**
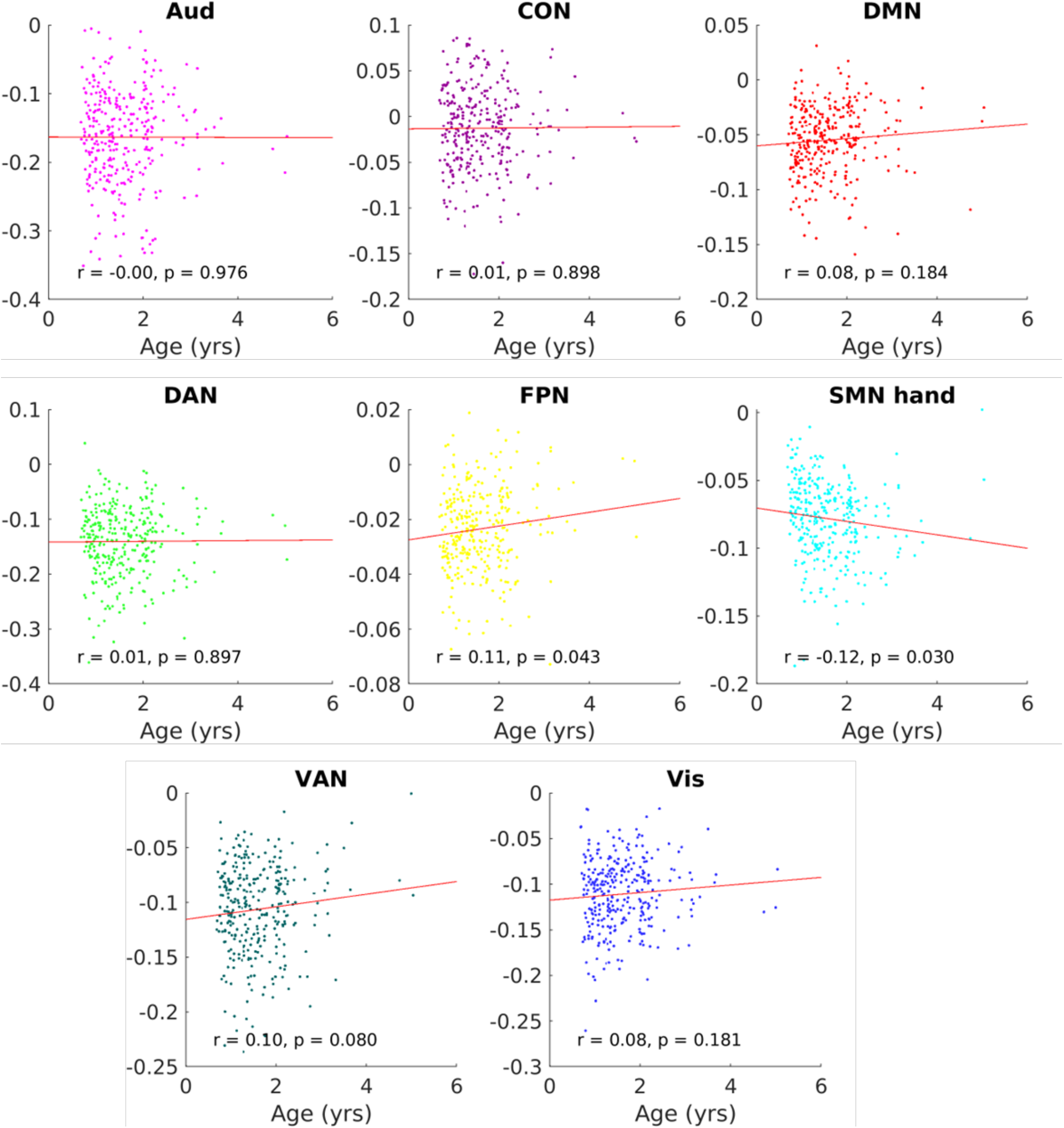
Within-network FC difference (All – Subset) across eight partially-retained networks.

**Supplementary Figure 10.**
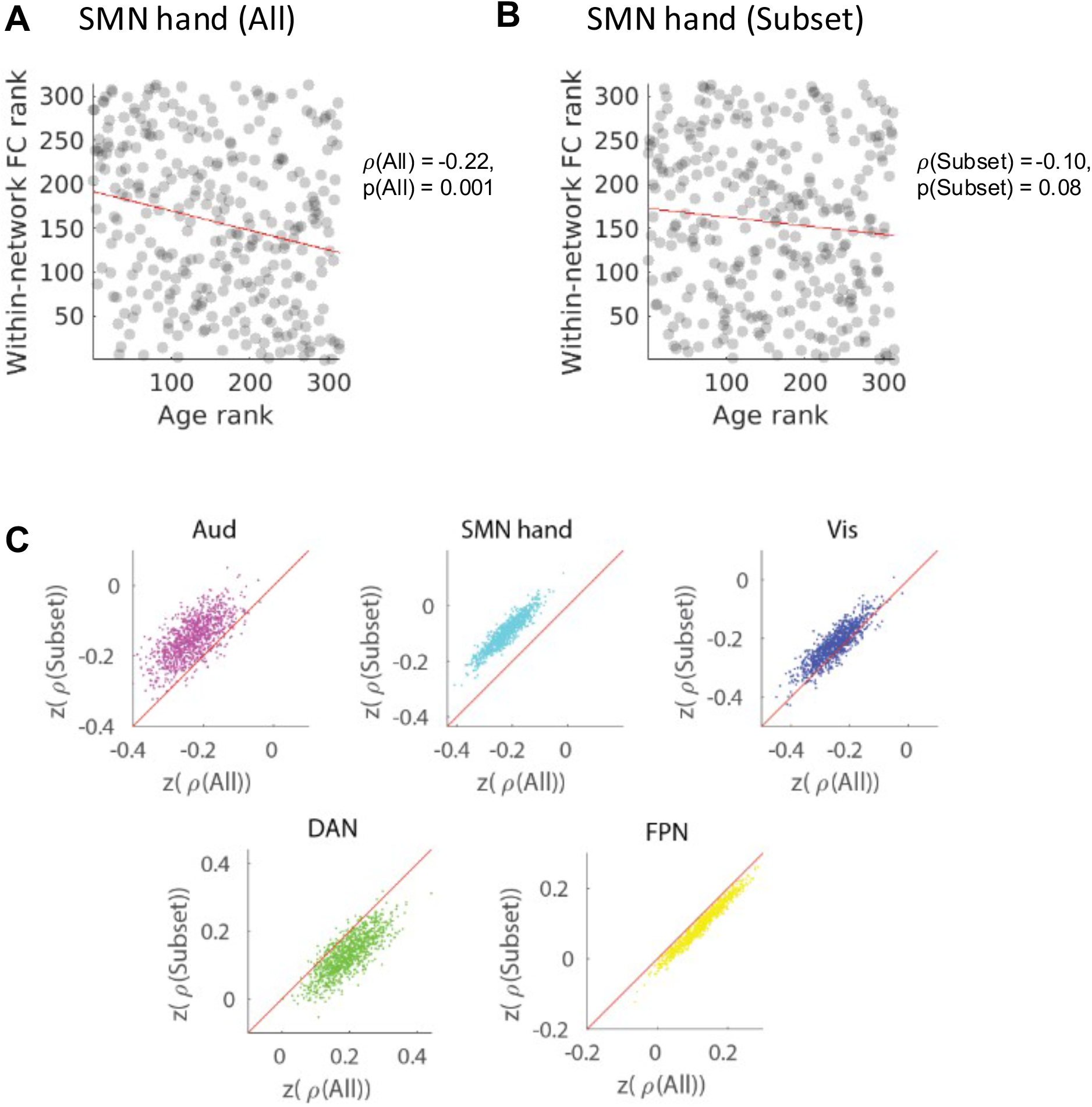
Correlation between age and within-network FC in using our area subset versus all areas. A) Scatter plot of within-network FC versus age for SMN hand network using all areas. B) Scatter plot of within-network FC versus age for SMN hand network using our area subset. C) The within-network FC for three networks is negatively correlated with age (Aud, SMN hand, Vis), and the within-network FC for two networks is positively correlated with age (DAN, FPN). The x-axis is the Fisher-Z-transformed Spearman’s correlation (*ρ*) between within-network FC using all areas and age. The y-axis is the Fisher-Z-transformed Spearman’s correlation (*ρ*) within-network FC using our area subset and age). Each data point presents a bootstrap sample of sessions (N = 1000). Red line shows the line of least-squared fit in A-B and the line of identity in C.

**Supplementary Figure 11.**
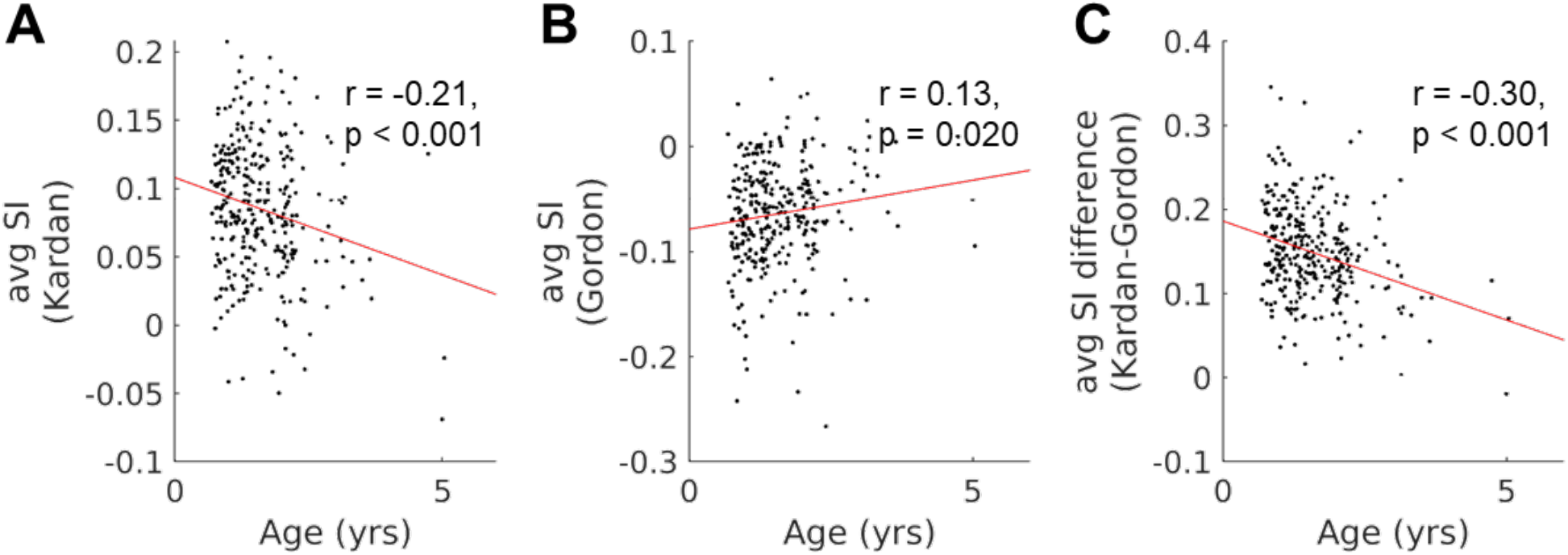
A scatter plot between chronological age in years and average SI for individual BCP sessions. A) infant networks (“Kardan”). B) adult networks (“Gordon”). C) Difference in infant networks (“Kardan”) and adult networks (“Gordon”).

**Supplementary Figure 12.**
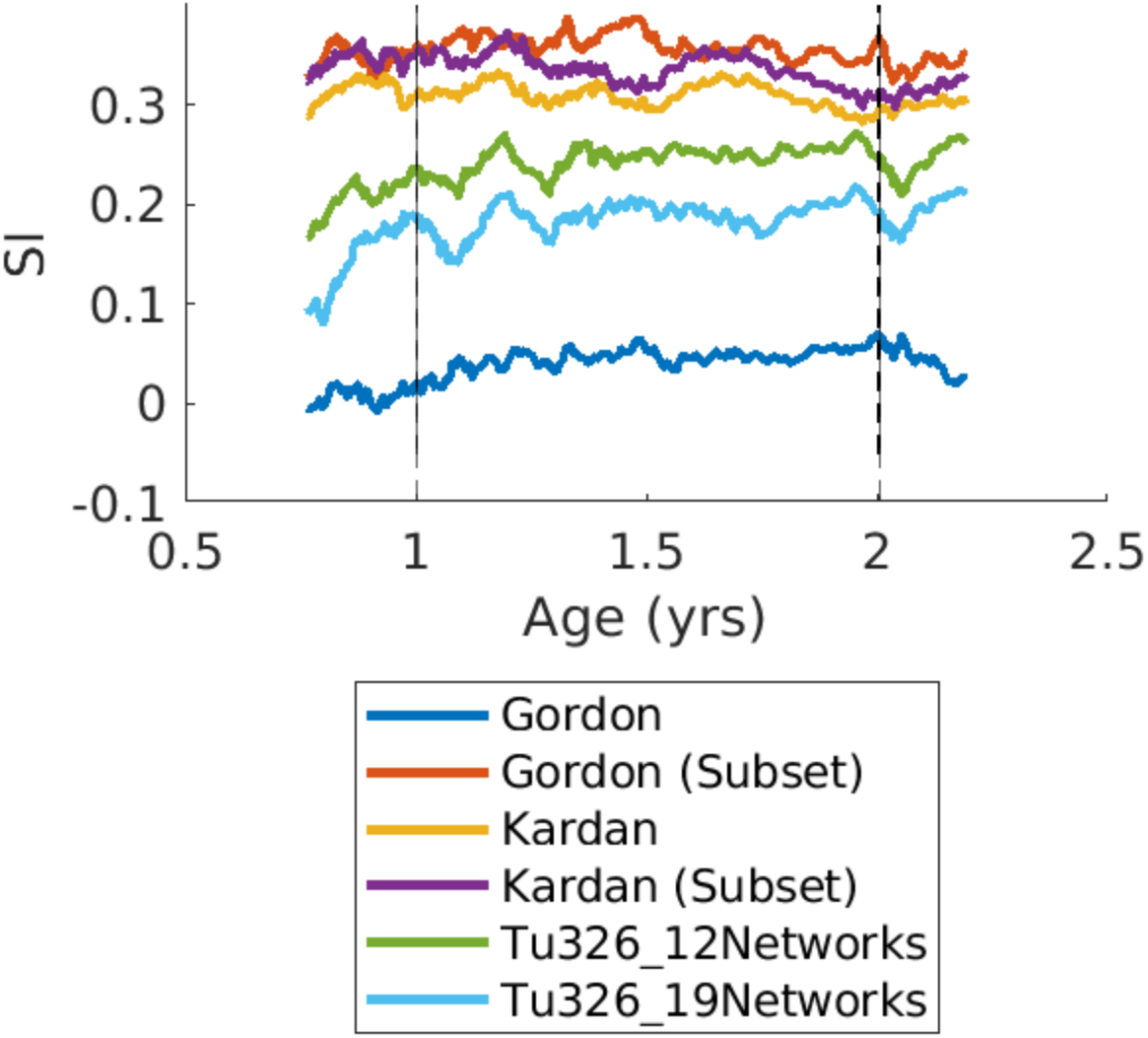
Moving average analysis (Figure 4A) adding the results using infant networks from an independent dataset.

**Supplementary Figure 13.**
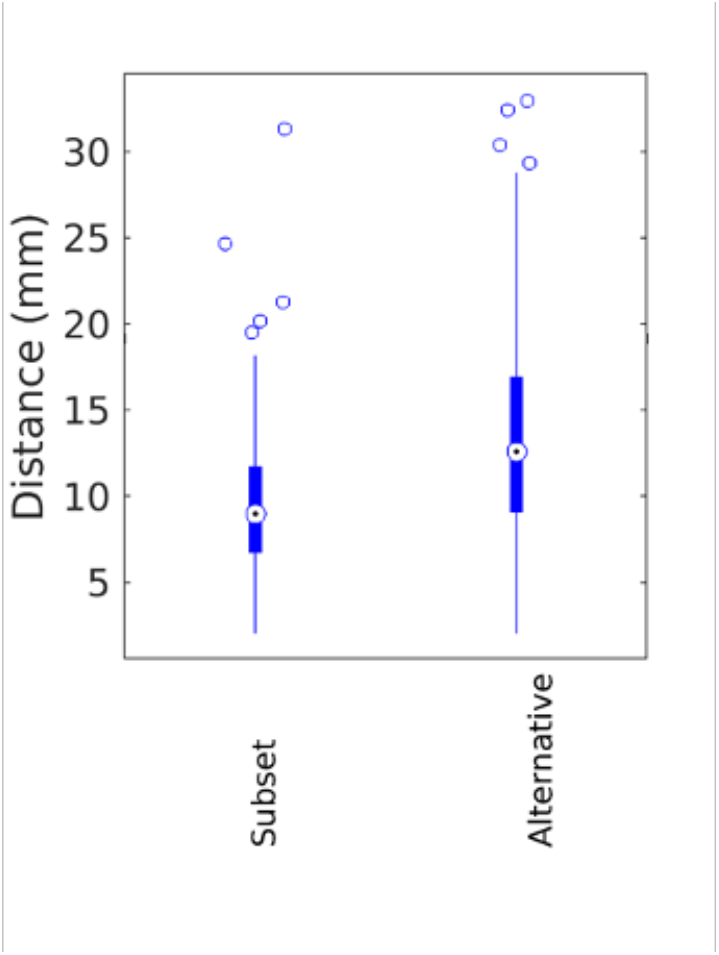
Distance between our area subset and alternative areas to the 153 Dworestky high consensus ROI.

**Supplementary Figure 14.**
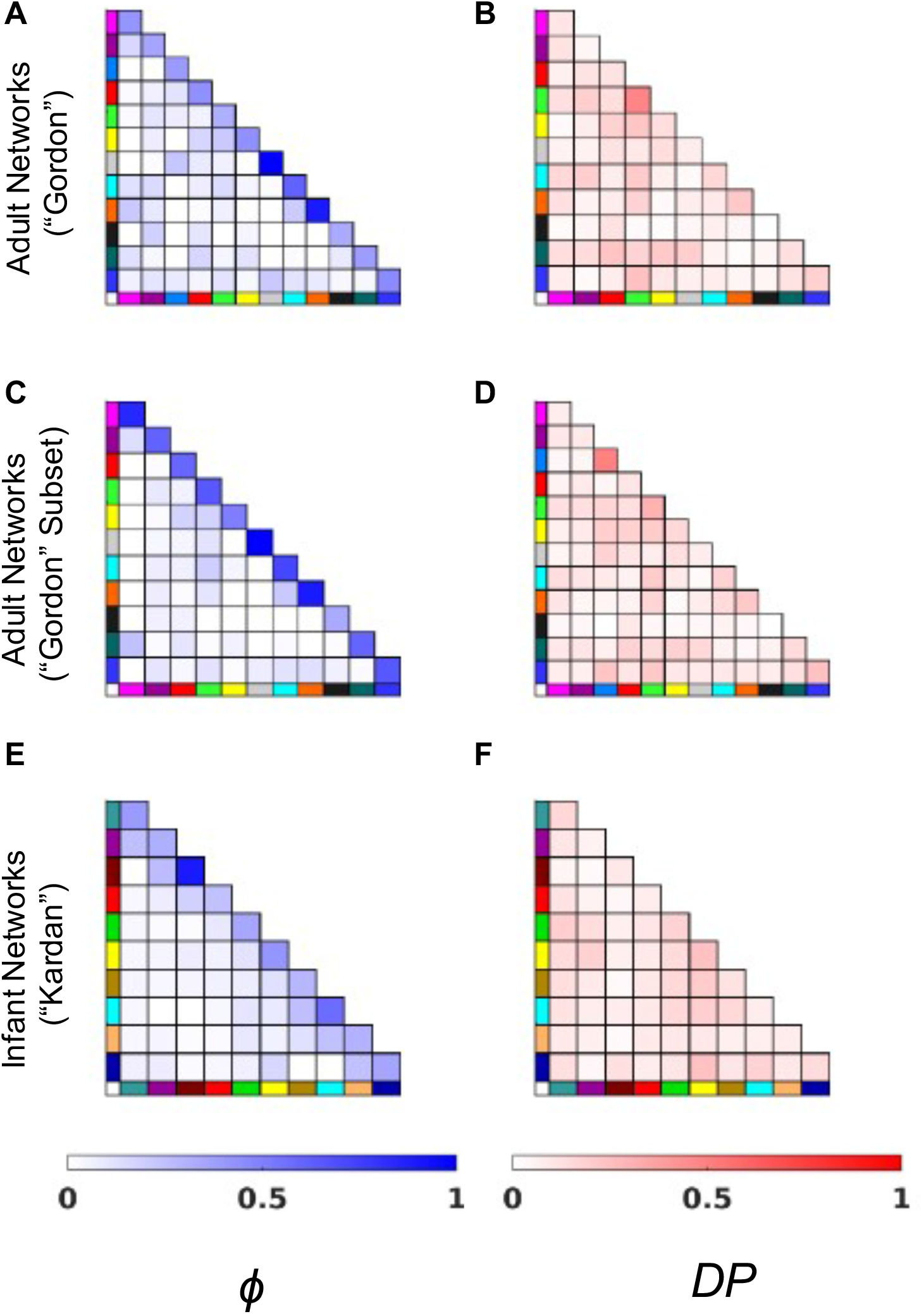
Fraction of high consistency (ϕ) and high differential power (DP) edges (top 10%) across (A-B) adult networks (“Gordon”), (C-D) adult networks (“Gordon” Subset), (E-F) infant networks (“Kardan”).

**Supplementary Figure 15.**
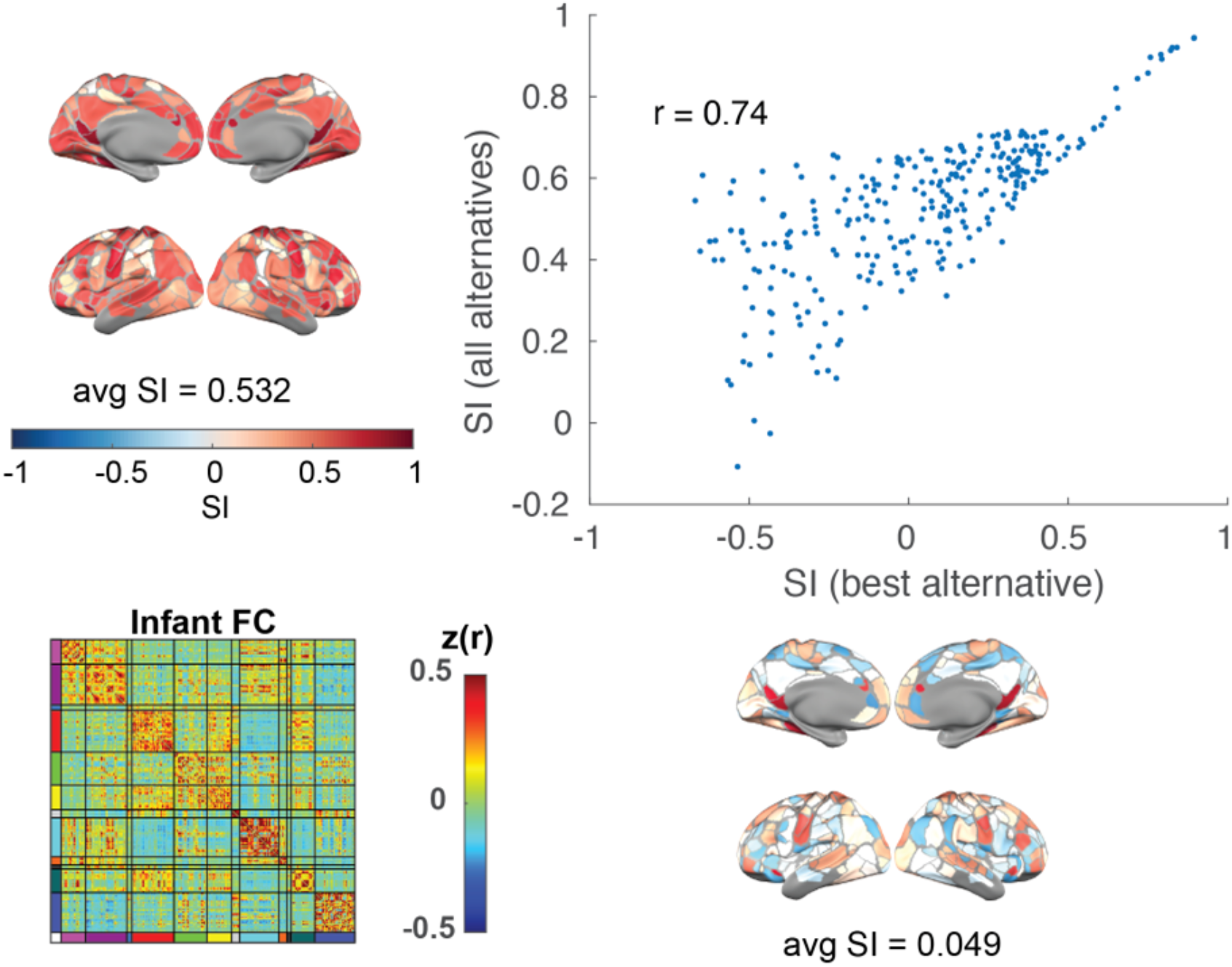
Correlation between silhouette index calculated with the best network with all alternative networks.

**Supplementary Table 1.**

| Supplementary Table 1 |  |  |  |
| --- | --- | --- | --- |
|  | Cohort | BCP | WashU 120 |
| Acquisition | Location | University of Minnesota | Washington University in St. Louis |
|  | Scanners | Siemens Prisma 3T Scanner | Siemens 3T Trio Tim |
|  | Headcoil | 32-channel | 12-channel |
|  | Sequence type | Gradient-echo EPI | Gradient-echo EPI |
|  | Resolution (BOLD) | 2mm isotropic | 4 mm isotropic |
|  | Phase encoding direction | AP+PA | AP |
|  | TR (s) | UMN - 0.72 (N = 70), 0.8 (N = 107) | 2.5 |
|  | TE (ms) | 37 | 27 |
|  | Resolution (T1) | 0.8 mm | 1 mm |
|  | Multi-band factor | 8 | N/A |
|  | State | Natural Sleep | Fixation on crosshair |
|  | Average clean data frames |  | 14.0 min |
|  | Frames per BOLD run | 420 | 120 |
|  | BOLD runs | 2-4 | 2 |
| Processing | Processing pipeline version | DCAN-Infant (0.0.22) | adult EPI (BOLD) preprocessing pipeline using the 4dfp tool suite |
|  | Distortion correction | ANTs SyN registration | None |
|  | Bias field correction | N4 method | None |
|  | Denoising | Respiratory notch filter, demean, detrend, 24 parameters nuisance regression, remove FD>0.3mm to apply bandpass filtering (0.008-0.09 Hz) then interpolate the missing frames | Demeaning and detrending, , multiple regression including: whole-brain, ventricular and white matter signals, and motion regressors derived by Volterra expansion (Friston et al. 1996), and a band-pass filter (0.009 Hz < f < 0.08Hz). |
|  | Scrubbing threshold | filtered FD<0.2mm, outlier (across-vertex STD on low FD frames >3MAD of the median of all frames) | FD<0.2mm, at least 5 consecutive low-motion frames |
|  | Tissue Segmentation | ANTs Joint Label Fusion was performed using a set of ALBERT atlases with segmentations generated and manually corrected by DCAN for ages 0-5 months old. For older ages, the atlases used for JLF were a set of 10 ABCD subjects for which we generated segmentations. Manual curation of tissue segmentation was performed where necessary. | FreeSurfer's default recon-all processing pipeline (version 5.0) |
|  | Surface reconstruction & registration | Modified FreeSurfer reconstruction (no hires, aseg from JLF, adjusted class means of tissue to fit T1w contrasts), registered to fs_LR32k using spherical registration (with MSMsulc). | FreeSurfer's default recon-all processing pipeline (version 5.0) |
| | BOLD data geodesic smoothing | $\sigma = 2.55$ mm | $\sigma = 2.55$ mm |
| **Quality Control** |  |  |  |
|  |  | BrainSwipes crowdsourcing ratings with average aggregated passing rate >75% for anatomical or functional images + manual screening. | N/A |
|  | Data availability | https://nda.nih.gov/edit_collection.html?id=2848 | https://legacy.openfmri.org/dataset/ds000243/ |

**Supplementary Table 2.**

| Supplementary Table 2 |  |  |
| --- | --- | --- |
| x | y | z |
| -18.8 | -48.7 | 65 |
| -51.8 | -7.8 | 38.5 |
| -18.4 | -85.5 | 21.6 |
| -47.2 | -58 | 30.8 |
| -38.1 | 48.8 | 10.5 |
| -55.9 | -47.7 | -9.3 |
| -14.4 | -57.8 | 18.4 |
| -8.8 | -49.8 | 4.2 |
| -11.3 | -83.2 | 3.9 |
| -1.7 | -17.7 | 39.1 |
| -10 | 33.9 | 21.5 |
| -10.7 | -47.5 | 60.3 |
| -15.6 | -33.1 | 66.1 |
| -10.9 | -29.3 | 69.5 |
| -6.6 | -20.4 | 74.2 |
| -10.8 | -41.1 | 64.9 |
| -5 | -28.2 | 60.4 |
| -5.4 | -15.9 | 48.8 |
| -35.8 | -29.7 | 54.5 |
| -41.5 | -12.5 | 50.4 |
| -42.1 | -4.5 | 47.3 |
| -27.3 | 1.9 | 52.9 |
| -19.8 | 6.4 | 55.7 |
| -19.5 | 30.1 | 45.5 |
| -36.8 | -22.8 | 61.9 |
| -20.5 | -24.9 | 64.5 |
| -23.4 | -13.8 | 64.2 |
| -17.2 | -8.6 | 67.9 |
| -28.6 | -44.7 | 61.7 |
| -31.1 | -48.9 | 47.1 |
| -42.9 | -45 | 43 |
| -51.5 | -11.9 | 29.7 |
| -51.7 | -30.9 | 39.9 |
| -27.5 | -37.2 | 61.4 |
| -47.2 | -31.4 | 54.8 |
| -46.1 | -17.8 | 52.7 |
| -44.8 | -54 | 14.6 |
| -51.6 | -55.9 | 11.4 |
| -48.1 | -40 | 2.4 |
| -46.3 | -41.4 | 25.9 |
| -52.7 | -20.6 | 5.4 |
| -58.7 | -29.9 | 11.1 |
| -40.6 | -38.3 | 14.5 |
| -38.7 | -16 | -5.3 |
| -50 | 20.8 | 10.6 |
| -37.7 | 2.9 | 11.7 |
| -40.3 | 50.4 | -4.8 |
| -32.5 | 17.2 | -7.8 |
| -44.3 | 33.2 | -7.2 |
| -45.4 | 28.8 | 0.8 |
| -20.4 | -64.6 | 51.4 |
| -34.1 | -61 | 42.4 |
| -31.3 | -84.2 | 9 |
| -34.2 | -86.6 | -0.5 |
| -46.2 | -57.7 | -7.9 |
| -55.1 | -32.3 | 23 |
| -43 | 19.4 | 33.5 |
| -40.2 | 23.6 | 23.3 |
| -48.6 | 7.5 | 11.1 |
| -5.9 | 54.8 | -11.3 |
| -6.8 | 38.2 | -9.4 |
| -33.8 | -33.2 | -15.4 |
| -28.8 | -58.8 | -9.1 |
| -34.4 | -63.9 | -15.7 |
| -34.3 | -43.8 | -21.6 |
| -5.4 | -88 | 18.6 |
| -8.6 | -77.5 | -3.5 |
| -22.6 | -81.7 | -11.7 |
| -22.5 | -37.1 | -15 |
| -15.9 | 48.6 | 37.2 |
| -19.5 | 56.3 | 27.5 |
| -21.3 | 63.1 | 1.9 |
| -28.6 | 50.9 | 10.1 |
| -6.5 | 54.7 | 18.1 |
| -15.7 | 64.7 | 13.7 |
| -26.2 | 26.6 | 38.8 |
| -29.3 | 16.8 | 50.7 |
| -41.7 | 16.1 | 47.5 |
| -54.4 | -1.4 | -0.7 |
| -59 | -18 | -3 |
| 20.8 | -48.2 | 66.1 |
| 49.6 | -7.4 | 36.1 |
| 22 | -84.6 | 23.7 |
| 47.9 | -42.5 | 41.5 |
| 38.1 | 45.9 | 7.7 |
| 59.7 | -41 | -10.9 |
| 13.8 | -54.1 | 10.9 |
| 15.5 | -74.1 | 9.4 |
| 6.7 | 5 | 55.9 |
| 8.4 | 34.7 | 22.6 |
| 3 | -19.6 | 37.9 |
| 8.8 | 10.8 | 45.9 |
| 16.5 | -32.8 | 67.7 |
| 4.8 | -27.1 | 64.8 |
| 11.9 | -40.7 | 67 |
| 5.1 | -17.1 | 51.6 |
| 6.8 | -8.1 | 50.9 |
| 42.3 | -11 | 47.3 |
| 42.5 | -2.3 | 47.2 |
| 29.2 | 1.9 | 52.4 |
| 21.9 | 21 | 46.2 |
| 38.1 | -22.4 | 60.3 |
| 19.7 | -25 | 65.2 |
| 12.4 | -28.3 | 69.6 |
| 29.2 | -13.5 | 64.2 |
| 17 | -16.9 | 70.9 |
| 20.9 | -6.4 | 65 |
| 29.5 | -42.5 | 60.4 |
| 38.8 | -42.6 | 40.4 |
| 53.9 | -8.3 | 26.1 |
| 28 | -34.8 | 63.1 |
| 39.2 | -34.6 | 57.5 |
| 37.3 | -25.9 | 50.9 |
| 47.8 | -15.1 | 49.3 |
| 48.9 | -53 | 28.6 |
| 57.5 | -45.3 | 9 |
| 60.9 | -38.7 | 1.7 |
| 54.9 | -27 | 29.6 |
| 57.1 | -17 | -2.6 |
| 53.8 | -15.8 | 5.2 |
| 47.4 | -39.6 | 13.2 |
| 45.5 | -37.3 | 3.4 |
| 48.5 | -26.5 | -0.1 |
| 60 | -25.2 | 10.2 |
| 38.8 | -14.4 | -5 |
| 42.8 | 48.3 | -5.1 |
| 45.2 | 30.7 | -5.6 |
| 30.6 | 22.8 | -4.7 |
| 7.7 | -85.6 | 31.6 |
| 35.4 | -77.1 | 21.1 |
| 41.5 | -53.5 | 44 |
| 33.5 | -48.2 | 49.4 |
| 31.7 | -85.7 | 2.4 |
| 43.8 | -67.2 | 2 |
| 57 | -53.8 | -1.1 |
| 37.8 | 28.7 | 35.6 |
| 41.8 | 29.1 | 21.6 |
| 50.1 | 3 | 3.9 |
| 38.6 | 18.8 | 25.5 |
| 28.4 | 57 | -5.1 |
| 4.8 | 65.1 | -7.1 |
| 7.2 | 48.4 | -10.1 |
| 34.6 | -35.6 | -12.3 |
| 34.6 | -23.9 | -20.4 |
| 26.9 | -69.1 | -6.6 |
| 34.9 | -44 | -20 |
| 13.8 | -92.3 | 14.7 |
| 10.5 | -73.8 | -1.5 |
| 20.4 | -87.3 | -6.6 |
| 5.1 | -80.2 | 23.1 |
| 24.5 | -36.2 | -13.2 |
| 21 | 32.8 | 42.1 |
| 21.4 | 42.8 | 35.1 |
| 23.5 | 59.1 | 4.9 |
| 30.9 | 52.2 | 9.9 |
| 8.2 | 53.8 | 14 |
| 5.9 | 54.9 | 29.4 |
| 13.8 | 46.7 | 42.1 |
| 6.8 | 44.5 | 34.8 |
| 30.6 | 18.9 | 48.7 |
| 42.4 | 19.5 | 48.2 |
| 38.9 | 9.6 | 42.7 |
| 39.7 | -22.5 | 2.6 |
| 55.8 | 2 | -2 |
| 57.1 | -6.3 | -7.7 |
| 46.6 | -21.5 | -8.5 |

**Supplementary Table 3.** Percentage of highly consistent (*ϕ*) edges across different network assignment schemes.

|  | Gordon All | Gordon Subset | Kardan |
| --- | --- | --- | --- |
| within-network | 44.1% | 63.1% | 49.5% |
| between-network | 8.3% | 6.5% | 5.4% |

**Supplementary Table 4.** Percentage of highly consistent (*ϕ*) edges across different network assignment schemes. The eight partially retained networks were bolded and had an asterisk.

| within-network | Gordon All | Gordon Subset |
| --- | --- | --- |
| **Aud\*** | 41.30% | 84.21% |
| **CON\*** | 35.64% | 61.73% |
| PMN | 40.00% | / |
| **DMN\*** | 43.17% | 59.07% |
| **DAN\*** | 31.45% | 65.17% |
| **FPN\*** | 43.12% | 50.84% |
| RTN | 100% | 100% |
| **SMN hand\*** | 61.45% | 72.82% |
| SMN mouth | 89.29% | 89.29% |
| Sal | 33.33% | 33.33% |
| **VAN\*** | 38.34% | 60.94% |
| **Vis\*** | 45.89% | 65.17% |

**Supplementary Table 5.** Percentage of high differential power (*DP*) edges across different network assignment schemes.

|  | Gordon All | Gordon Subset | Kardan |
| --- | --- | --- | --- |
| within-network | 16.50% | 14.73% | 12.79% |
| between-network | 10.81% | 10.98% | 9.93% |

**Supplementary Table 6.** Percentage of high differential power (*DP*) edges across different network assignment schemes. The eight partially retained networks were bolded and had an asterisk.

| **within-network** | **Gordon All** | **Gordon Subset** |
| --- | --- | --- |
| **Aud\*** | 7.97% | 10.53% |
| **CON\*** | 8.33% | 4.94% |
| PMN | 50.00% | / |
| **DMN\*** | 10.73% | 11.59% |
| **DAN\*** | 30.24% | 47.19% |
| **FPN\*** | 15.58% | 13.91% |
| RTN | 7.14% | 7.14% |
| **SMN hand\*** | 17.92% | 15.82% |
| SMN mouth | 21.43% | 21.43% |
| Sal | 0% | 0% |
| **VAN\*** | 14.62% | 12.02% |
| **Vis\*** | 24.97% | 18.43% |

